# Changing environments and genetic variation: inbreeding does not compromise short-term physiological responses

**DOI:** 10.1101/520015

**Authors:** James Buckley, Rónán Daly, Christina Cobbold, Karl Burgess, Barbara K. Mable

## Abstract

- Selfing plant lineages are surprisingly widespread and successful in a broad range of environments, despite showing reduced genetic diversity, which is predicted to reduce long-term evolutionary potential. However, short-term capacity to respond appropriately to new conditions might not require high levels of standing genetic variation. The purpose of this study was to directly test whether mating system variation and its associated changes in genetic variability in natural populations affected responses to short-term environmental challenges.
- We compared relative fitness and metabolome profiles of naturally outbreeding (genetically diverse) and inbreeding (genetically depauperate) populations of a long-lived perennial plant, *Arabidopsis lyrata,* under constant growth chamber conditions and an outdoor common garden environment outside its native range.
- We found no effect of mating system on survival or reproductive output, although several phenological traits showed different associations with latitude for outcrossing and inbreeding populations. Natural inbreeding had no effect on the plasticity of physiological responses, using either multivariate approaches or analysis of variation in individual metabolites. Moreover, while both growing environment and time significantly affected the relative abundance of individual metabolites, inbreeding populations responded similarly to outbreeding populations, suggesting adaptation to the outdoor environment, regardless of mating system.
- We conclude that low genetic diversity in naturally inbred populations may not compromise fitness or short-term capacity for appropriate physiological responses to environmental change. The absence of natural costs of inbreeding could help to explain the global success of clonal or asexual mating strategies for adapting to a wide range of environments.

## INTRODUCTION

Genetically informed conservation management programmes often assume that adaptive potential is limited by the amount of additive genetic variation maintained in a population (O’Brien, 1994; Hoffmann, Sgro, and Kristensen, 2017). However, many geographically widespread and invasive plant species are self-fertilising or asexually reproducing and the evolutionary transition from an outcrossing to a selfing mating system has occurred frequently (e.g. Igic, Lande, and Kohn, 2008; Razanajatovo et al., 2016). Such shifts are associated with increased genome-wide homozygosity, as well as reduced efficacy of purifying and positive selection (e.g. Wright, Kalisz, and Slotte, 2013). Inbreeding is thus thought to compromise long-term evolutionary potential through erosion of genetic variation and increase the risk of extinction through inbreeding depression, defined as the exposure of deleterious recessive mutations through increased homozygosity (Charlesworth and Charlesworth, 1987; Keller and Waller, 2002). Consistent with this, selfing lineages are predicted to show higher extinction rates than related self-incompatible lineages (Goldberg et al., 2010). Together, these lines of evidence support the view that the long-term potential to adapt to environmental change will be compromised in low diversity selfing (inbred) lineages, compared to higher diversity outcrossed lineages (Bijlsma and Loeschcke, 2012; Wright, Kalisz, and Slotte, 2013).

However, multiple lines of evidence suggest that low levels of genetic variation associated with selfing may not affect the ability of lineages to respond appropriately to shorter-term environmental change. Firstly, many highly self-fertilising or asexually reproducing plants (modes of reproduction that reduce genetic variation) show broad distributions, and are able to invade and colonise new locations (Barrett, Colautti, and Eckert, 2008; van Kleunen et al., 2008; Razanajatovo et al., 2016). There are multiple examples of self-fertilising species rapidly colonising and adapting to new environments (Colautti et al., 2017; Willoughby et al., 2018), despite starting with low levels of genetic diversity. Secondly, low levels of neutral genetic variation may not always equate to low levels of adaptive genetic variation, and therefore may be a poor proxy for the evolutionary potential of a population (Whitlock, 2014). The loss of individual selfed or clonal lineages with a high genetic load can also reduce the impacts of inbreeding depression at the population-level (Keller and Waller, 2002; Hedrick and Garcia-Dorado, 2016). Finally, selfing has obvious advantages when reproducing in a new environment where conspecifics are scarce (Eckert et al., 2010), but few studies have directly tested the effects of the resulting low additive genetic variation on short-term physiological plasticity.

Experimental laboratory studies suggests that the negative effects of inbreeding on trait plasticity may be strongest under stressful environments (Bijlsma and Loeschcke, 2012). For example, artificially inbred families or experimental lines show reduced survival under extreme temperature stress (Kristensen et al., 2008), reduced tolerance to herbivores (Ivey, Carr, and Eubanks, 2004), and reduced induction of anti-predator or anti-herbivore defense traits (Auld and Relyea, 2010; Campbell, Thaler, and Kessler, 2012; Kariyat et al., 2012; Campbell et al., 2014; Swillen, Vanoverbeke, and De Meester, 2015). However, in other experiments, the effects of inbreeding on trait plasticity were either not observed (Schlichtling and Levin, 1986), or varied among traits and were not consistent across inbred families (Ivey, Carr, and Eubanks, 2004; Schou, Kristensen, and Loeschcke, 2015). So, even in a context of experimental inbreeding in normally outcrossing species (when inbreeding depression should be high), short-term responses might not always be compromised by reduced heterozygosity or diversity in inbred lineages.

Recent work on the molecular basis of inbreeding effects on plasticity has revealed altered gene expression patterns associated with artificially inbred lines, as well as interactive effects of environmental stress and inbreeding on gene expression (Kristensen et al., 2010; Paige, 2010). Inbred *Drosophila* lines exposed to temperature stress showed increased variance in metabolite profiles relative to outcrossed lines, as well as consistent effects of inbreeding on particular metabolites (Pedersen et al., 2008). Similarly, experimentally inbred plant families (*Solanum carolinense*) showed reduced expression of anti-herbivore defensive metabolites (Campbell, Thaler, and Kessler, 2012). Thus, it seems clear that an initial shift from outcrossing to inbreeding could compromise appropriate metabolic responses to stressors through inbreeding depression. However, most studies have compared the consequences of artificially-induced inbreeding, or mating system variation between species, rather than within species. We know far less about how populations with a sufficiently long history of inbreeding to purge deleterious recessive mutations will be able to adapt to changing environmental conditions, despite reduced levels of genetic variation compared to their outcrossing relatives.

To test how natural mating system variation within a species impacts physiological responses to abrupt environmental change we used *Arabidopsis lyrata*, a predominantly outcrossing perennial herb that shows natural variation in mating system around the Great Lakes region in North America (Mable et al., 2005; Mable and Adam, 2007; Foxe et al., 2010). Inbreeding populations, in which the majority of individuals are capable of self-fertilisation, show significantly reduced heterozygosity and genetic diversity relative to outcrossing populations (Foxe et al., 2010; Buckley et al., 2016), although they show only minor changes in floral morphology consistent with the evolution of a selfing phenotype (Carleial, van Kleunen, and Stift, 2017b). Populations around the Great Lakes occupy several distinct habitats (rocky alvar and sand dune) and are distributed across a broad latitudinal gradient, characterised by variation in climate and the length of the growing period, so are also an interesting system in which to study local adaptation and its interaction with mating system. Patterns of population genetic structure suggest that the loss of self-incompatibility arose multiple times during several independent postglacial colonisations of the North American Great Lakes region (Hoebe, 2009; Foxe et al., 2010).

Experimentally-induced inbreeding in outcrossing *A. lyrata* populations from Europe has been found to result in strong inbreeding depression in growth and germination-related traits, as well as changes to constitutive patterns of gene expression under stable environmental conditions (Sletvold et al., 2013; Stift et al., 2013; Menzel et al., 2015). In contrast, experimentally-inbred North American outcrossing populations show more subtle fitness reductions (Stift et al., 2013) and show suprisingly few differences compared to geographically proximate inbreeding populations. While naturally inbreeding populations from North America showed significantly reduced germination rates compared to closely-related outcrossing populations, no inbreeding depression was found for seedling growth rates or induced defense responses when challenged with herbivores for either mating system type (Joschinski, van Kleunen, and Stift, 2015; Carleial, van Kleunen, and Stift, 2017a). Moreover, inbreeding load has been found to be low and not substantially different in naturally inbreeding and outcrossing populations from North America, although the former show a greater increase in fitness when crossed to other populations (Willi, 2013). Similarly, in common garden experiments using experimental crosses between populations, heterosis was found to be higher in inbred compared to outcrossed populations, but magnitudes of inbreeding depression were similar regardless of mating system (Oakley, Spoelhof, and Schemske, 2015). North American populations of *A. lyrata* show a substantial reduction in genetic diversity compared to European populations, suggestive of a historical bottleneck (Ross-Ibarra et al., 2008; Mattila et al., 2017), but also that there has been some purging of the genetic load in both outcrossing and inbreeding populations (Stift et al., 2013; Willi, 2013). *Arabidopsis lyrata* is therefore a good model to assess the impacts on adaptive potential of loss of genetic diversity within a species caused by inbreeding without being overwhelmed by large differences in inbreeding depression in relation to mating system. Previous physiological studies in *A. lyrata* have revealed extensive variation in metabolite profiles among populations from different geographic regions (Davey et al., 2008; Kunin et al., 2009), as well as divergence in cold tolerance responses among regions (Davey, Woodward, and Quick, 2008), but these analyses were restricted to outcrossing populations from Europe. An important gap in our knowledge is thus whether natural variation in levels of inbreeding and genetic diversity affects short-term physiological responses to environmental change.

The purpose of this study was to test whether naturally inbred populations show reduced fitness and altered physiological responses in a common garden environment when compared to outbred populations. The common garden environment was situated outside the native range of *A. lyrata* and therefore provided growing conditions that differed from those naturally experienced. Specifically, we asked: 1) Is inbreeding associated with reduced fitness compared to outcrossing populations when individuals are transplanted to the common garden environment? 2) Is there a change in the metabolome over time when plants are transplanted to a naturally variable environment compared to those that are kept under constant environmental conditions? 3) Does inbreeding alter the direction or magnitude of physiological plasticity over time or across environments?

Our results reveal no consistent effects of mating system on fitness or short-term physiological responses to environmental changes, despite clear metabolomic divergence under the two growing environments at the later time point. The remarkable similarity in metabolic responses of inbred and outcrossed populations suggests that standing genetic variation is not important for adaptive physiological plasticity under potentially stressful novel environments.

## MATERIALS AND METHODS

### Seed sampling and plant origins

Seeds were collected from 25-40 individual *A. lyrata* ssp. *lyrata* plants per population from thirteen sites across the North American Great Lakes region (Fig 1a; Table S1), with eight outcrossing and five inbreeding populations selected based on a combination of outcrossing rates (*t_m_*) estimated using progeny arrays, proportion of self-compatible individuals (reflecting potential for inbreeding), and observed heterozygosity (*H_o_*; reflecting actual history of inbreeding) in a previous study (Foxe et al., 2010)(see Table S2 for values of *t_m_* and *H_o_*). Seeds were collected in 2011 (Table S2), except for two inbreeding populations (KTT in 2007 and PTP was supplemented in 2012). Outcrossing populations have significantly higher observed heterozygosity and nucleotide diversity relative to inbreeding populations based on both neutral microsatellite markers (Table S2; Foxe et al. 2010) and RAD-seq markers (Buckley *et al.* 2016). A mixed mating population TSSA *(t_m_* = 0.41) was grouped with the outcrossing populations (*t_m_* = 0.83-0.99) rather than the inbreeding populations (*t_m_* = 0.09-0.31), as it showed similar levels of genome-wide diversity and heterozygosity to the outcrossing populations (Buckley et al., 2016). Most populations were found in freshwater coastal sand dune habitats, though plants at some sites (TC and TSSA) occupied rocky, alvar habitats and KTT was found in an oak woodlands sand flat. The inbreeding populations are predicted to represent three putatively independent origins of selfing (Foxe et al., 2010), with geographically proximate outcrossing populations predicted to be in the same genetic cluster included, where possible for metabolomics analysis: 1) inbreeding: RON, PTP, LPT; outcrossing: PCR, PIN; 2) inbreeding: TC; outcrossing TSS, TSSA; 3) inbreeding KTT is not located on a lakefront but it is on a similar latitude as IND, which is in the same genetic cluster as SAK, SBD, and MAN (Table S1).

**Figure 1:**
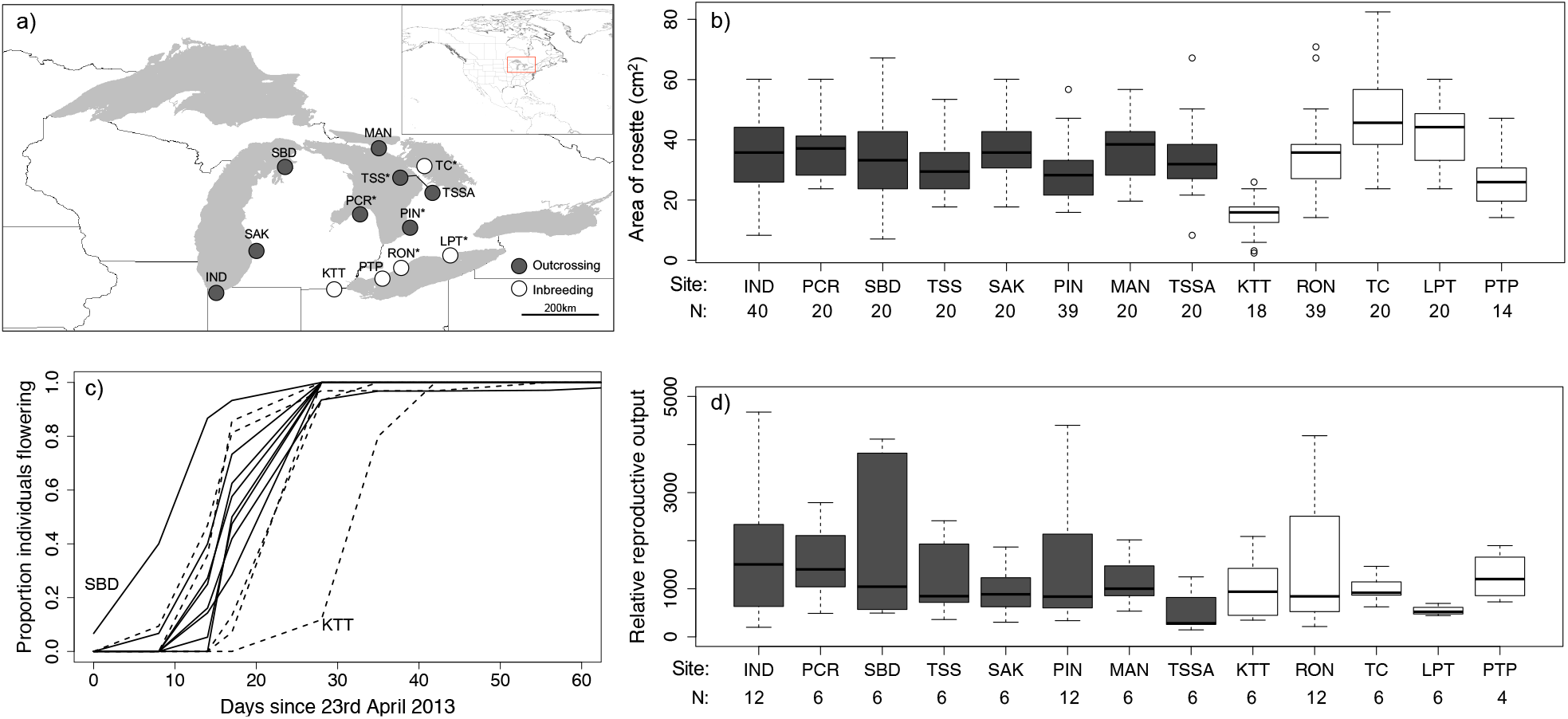
*Arabidopsis lyrata* sampling location and variation in phenology and fitness among populations and with respect to population mating system, (a) map of sampling sites with eight outcrossing and five inbreeding populations and an * indicating those populations used for metabolomics analysis; (b) boxplot of rosette area (mm^2^) before transplantation to common garden; (c) proportion of plants flowering per population at 8 time points over an 8-week period (dashed lines = inbreeding populations; solid lines = outcrossing populations; (d) boxplot of reproductive effort (average length of fruits multiplied by number of fruits produced) on 1^st^ July 2013. For (a), (b) and (d), dark grey indicates outcrossing populations, and white indicates inbreeding populations. Site codes and sample sizes are given under plots (b) and (d), in which populations are ordered by decreasing outcrossing rates (from left to right).

### The effects of population inbreeding on growth, survival and reproduction

#### Experimental Design

To compare relative fitness of outbred and inbred *A. l. lyrata.* we established a common garden in a novel environment in Scotland (at the University of Glasgow Scottish Centre for Ecology and the Natural Environment, SCENE, on Loch Lomond; 56.1289°N, 4.6129° W). The summer months in this part of Scotland tend to be relatively cool and wet, and winter months more mild, than the corresponding times of the year around the North American Great lakes, where hot and dry summers and cold winters are more common. The common garden site was situated in a clearance surrounded by deciduous woodland.

We therefore expected this common garden to represent a novel environment for *A. lyrata.* Seeds were germinated under controlled growth chamber conditions (16h: 8h, 20°C:16°C, light: dark cycle) in Levington F2+S (seed & modular + sand) compost in 40-cell seedling propagators (Parasene, Cradley Heath, UK) to maximise germination rates. Three seeds from each of 20 maternal families per population were germinated per tray cell and thinned to one seedling per maternal family per population. For three populations (outcrossing IND and PIN and inbreeding RON) we used one seedling from each of 40 maternal seed families to obtain more precise estimates of fitness. These were three of the largest populations of *A. lyrata* in the field, and therefore were sites from which seeds from a greater number of maternal families were available. For one inbreeding population (PTP) only 14 maternal seed families were available (Table S2).

In total, 310 individuals, 199 of which were from outcrossing populations and 111 from inbreeding populations, were transplanted to the common garden on 21^st^ September 2012. They were placed in a raised bed filled with Levington F2+S compost and horticultural grit to enhance drainage, and surrounded by high wire fencing to exclude mammalian herbivores. Two iButton dataloggers were placed in the centre of the plot to record above-ground and below-ground temperatures. Four propagator trays with 5-10 individuals from each population formed a block and four blocks were placed in a linear common garden plot (Fig S1). Plants in the common garden were watered once after transplanting, but thereafter no additional water or fertiliser was added during the experimental period. Seedlings from different populations were systematically arranged across the trays to distribute populations differing in mating system evenly across the common garden plot (Fig S1). Rosette size on transplant (7 weeks after germination) was calculated as the area of a circle from two perpendicular measurements of rosette diameter.

#### Survival and reproductive phenology in the common garden

Plants were checked approximately once a week in the spring from 23^rd^ April to 4^th^ June (and then every 2 weeks till 9^th^ July) to record the proportion of plants with at least one open flower. All plants, except two individuals had flowered by early June. At the beginning of July, we counted the total number of fruits produced by two haphazardly selected individuals per population from each of three experimental blocks (in total 6 individuals per population, or 12 for populations IND, PIN and RON), to provide a snapshot measure of reproductive effort. Mean fruit length, which is highly correlated with seed number (Hoebe, 2009), was measured, on average, from five haphazardly selected, fully-developed fruits from each individual. Relative reproductive output per individual for this subset was then calculated as the number of fruits multiplied by the mean fruit length. The proportion of plants surviving overwinter was recorded in late spring on 28^th^ May 2013 and 21^st^ May 2014. Survival in 2014 therefore represents cumulative fitness over the time course of the experiment.

#### Statistical analyses

The effect of population mating system on each response variable (plant rosette area before transplantation, proportion plants flowering, survivorship, number of fruits, mean length of fruits per plant and the combined measure of relative reproductive output) was tested using generalised linear mixed effects models (GLMMs) using the R package *lme4* (Bates *et al.* 2015). Population mating system was included as a fixed factor in the models, and population was modelled as a random effect, to account for unexplained variation due to geographic source. For traits where all plants were measured (rosette area, proportion flowering, survivorship) experimental block was also included as a separate random factor in this model. Where traits were only measured for a subset of individuals (fruit length, number fruits, reproductive output) there were insufficient individuals to estimate random effects of experimental block. To account for multiple testing, a Bonferroni-correct P-value threshold of 0.008 (0.05 divided by 6) was used to identify significant mating system effects. Binomial error distributions were used for modeling binary traits (survival and flowering status) and a normal error distribution used for continuous traits (rosette area and reproductive output). When modeling survival and propensity to flower, rosette size at transplant was included as a covariate to account for differences in initial growth rates in the cabinets. Model residuals were inspected to ensure good model fit and where necessary the response variables were log-transformed. To directly test for differences among populations in each response variable, we used separate models for a fixed effect of population on the different response variables controlling for the random effect of experimental block, but not considering mating system. Again a reduced P-value threshold of 0.008 was used to account for multiple testing and identify significant relationships. As multiple fruits were measured per individual, we additionally used a linear mixed effects model to estimate the proportion of variance in fruit length associated with population, maternal plant (family) and within-plant (residual variance). Likelihood ratio tests were used to estimate the significance of fixed effects following their removal from a model.

We additionally tested for variation in fitness-related traits consistent with local adaptation along the latitudinal range (41.6206 −45.6703 decimal degrees) occupied by these populations, and whether any of the fitness-related traits varied with population mating system. We conducted regression analyses to test for a significant interaction between mating system (inbreeding and outcrossing) and latitude for each of the above-described fitness-related traits, as well as the separate effect of latitude. The statistical models used for each trait were the same as described above.

### The effects of population inbreeding on the metabolome

#### Experimental Design

To determine whether population mating system affected the physiological response under contrasting environmental conditions, we compared metabolomic profiles over time for three outcrossing (PIN, PCR and TSS) and three inbreeding (RON, TC, LPT) populations when transplanted outdoors to the common garden, or when kept in a controlled growth chamber. We selected the three outcrossing and three inbreeding populations to represent similar levels of genetic structure and geographic range within each mating system group. Specifically, previous research showed that the chosen populations represent two different genetic clusters, with PIN, PCR, RON and LPT in one and TSS and TC in another (Foxe *et al.* 2010; Buckley *et al.* 2016). We germinated seeds from five maternal plants from each population under controlled growth cabinet conditions for six weeks. In August 2013 one seedling from each mother was either transplanted to new trays (6cm deep) under the same growth chamber conditions, or transplanted to the common garden environment (used in the fitness experiment).

#### Sampling strategy and methods for collecting leaf tissue

Two similar-sized leaves were sampled from the rosette of each individual at three time points in the common garden environment to examine changes in the metabolome over time. Each individual was sampled: 1) before transplanting to the outdoor environment (4th August 2013) to establish baseline profiles; 2) ~24h after transplantation to test for effects of ‘transplant shock’ (7th August 2013); and 3) 1 month after transplantation (5th September 2013), to give time for the plants to physiologically adapt to the growing environment. Any dirt was quickly removed from sampled leaves, before leaves were placed in a cryo-tube and flash frozen in liquid nitrogen. They were kept frozen on dry ice for a maximum of 2h during transport and then transferred to a −70°C freezer. The samples from the growth chamber plants were collected on the same days and in the same manner as described above. The plants were checked for signs of disease or herbivore damage and photos were taken of trays just before sampling to monitor their general status (e.g. colour of leaves).

#### Metabolomic data generation

For the three inbreeding and three outcrossing populations, individuals were selected for metabolite screening if leaf samples were available at all three time points for the two experimental growing conditions. Individuals from the same maternal families were used in the common garden experiment and the growth chamber. To allow us to assess changes in the metabolome over three timepoints and two treatments, metabolomics data was generated for only three individuals per population (nine individuals per mating system group). We therefore had insufficient power to resolve population-level differences in metabolomic plasticity through time and under different growing environments. One leaf per individual was placed in a Fastprep RNA bead tube (Lysing matrix D) and disrupted thoroughly in a Fastprep machine (MP Biomedicals). Each tube was dipped in liquid Nitrogen, and then disrupted in 2 × 10s runs, with samples refrozen in between runs. Then 1mL of chilled (−15°C) extraction buffer (chloroform: methanol: water in a 1:3:1 ratio) was added and the tube vortexed for 5s. The mixture was shaken on ice for 10min and centrifuged for 30s to separate tissue and lysate. The clear lysate was transferred to a new tube and used for metabolomics analysis. Briefly, 10μL of each sample was introduced to a liquid chromatography system (UltiMate 3000 RSLC, Thermo, UK) and separated on a 4.6 mm × 150 mm ZIC-pHILIC analytical column with a 2 mm × 20 mm guard column. The eluents were A: water with 20 mM ammonium carbonate and B: acetonitrile. The gradient ran from 20% A, 80% B to 80% A, 20% B in 15min with a wash at 95% A for 3min followed by equilibration at 20% A for 8min. Metabolites were detected using an Orbitrap Exactive (Thermofisher, UK) instrument in positive/negative switching mode at resolution 50,000 with a m/z scan range of 70-1400. In total, 108 samples, plus a sample of pooled individual extractions for quality control, were run in a randomised order interspersed with twelve blank extraction buffer samples. The LC–MS data were annotated using a bespoke bioinformatics pipeline (mzMatch, IDEOM and PiMP) developed at Glasgow Polyomics (Scheltema et al., 2011; Creek et al., 2012; Gloaguen et al., 2017). IDEOM assigns confidence scores ranging from 0-10 to each putative compound identification. We considered all compounds with an IDEOM score of at least 6 that were present at either the first or third experimental time point, which resulted in a final dataset of 936 metabolites. The raw peak heights for each putative compound in each individual sample was then corrected by subtracting the average of the twelve blank readings for that identified compound. These corrected peak heights were used for further analysis. The identity of 106 metabolites was confirmed through comparison of retention times and masses to a panel of 122 standards, of which 96 compounds were uniquely identified and 13 had two related identifications (from now referred to as ‘confidently identified metabolites’).

### Metabolomics Analyses

#### Multivariate Analyses

Principal Components Analysis (PCA) was performed on the dataset of 936 metabolites measured over the three time points using the R function *‘prcomp’*. We plotted the first two principal components against each other to explore broad patterns of change in the metabolite data with respect to experimental growing condition, time point and population mating system. As a measure of plasticity, we compared the magnitude and direction of metabolome shifts in response to the two different growth conditions (growth chamber and outdoor common garden). We also plotted the difference in values of the first five principal components (PCs) for related individuals (same maternal family) growing in the different environments at time point 3 as a measure of relative plasticity. If reduced genetic variation from multiple generations of selfing compromises plasticity in key traits, we predicted that individuals from inbreeding populations would show a reduced magnitude of change in each PC relative to those from outcrossing populations. To test this, we used a two-way ANOVA with mating system as a fixed effect and the magnitude of this difference for each of the first five PCs separately as a response variable.

#### Metabolite diversity

Metabolite diversity was also estimated using a set of diversity measures that have been developed to assess the relative importance of differences in the abundance of species, as well as the presence or absence of species in a community (Leinster and Cobbold, 2012). Specifically, the emphasis placed on relative abundance is changed by adjusting a parameter q. In the context of metabolite data, q = 0 is equivalent to the total number of metabolites observed (with all metabolites weighted equally), whereas q = 1 or higher results in lower abundance metabolites having less influence on the diversity measures (i.e. the most abundant compounds shape diversity estimates). Here, we should note that using an untargeted metabolomics approach means that variation in raw peak heights among identified metabolites may reflect both inherent differences in metabolite detectability, as well as variation in their actual abundance. Estimating diversity using q = 1 therefore assumes that variation in metabolite detectability does not vary over orders of magnitude, which may not always be the case. Nevertheless, our ability to estimate the ‘true’ diversity of metabolites should not impact our statistical comparison of changes in diversity profiles over time in each growing environment for inbred and outcrossed individuals. We therefore statistically tested whether variation at q = 0 (metabolite “richness”) and q = 1 (when relative abundance is considered) was explained by time, environment, mating system or their interactions using GLMMs, with likelihood ratio tests used to determine the significance of individual terms and population included as a random effect. Non-significant interactions and individual terms were sequentially removed until a reduced final model consisting of only significant interactions and terms remained.

#### Changes in individual metabolites and metabolite pathways

To comprehensively test for the importance of mating system in explaining variation in metabolite concentrations, we used GLMMs to model variation in corrected peak heights for each of the 936 metabolites. We accounted for the random effect of population and tested the fixed effects of mating system, time, treatment, and all two-way and three-way interaction. Given the difficulty of interpreting three-way interactions, we also tested the effects of time, mating system and their interaction for each experimental condition separately. Given that time points 1 and 2 showed similar multivariate metabolomic patterns, we focused on data from time points 1 and 3 in the analysis for each environment separately. We corrected for multiple testing using the Benjamini-Hochberg procedure for restricting the false discovery rate to 5%. For the subset of confidently-identified metabolites, we identified those metabolites that were on average > 1-fold higher or lower in the common garden samples relative to the growth chamber samples at time point 3, but which showed no difference (< 1-fold changes) at time point 1 (when all plants were in the growth chamber). For these confidently identified compounds we also tested for a significant interaction between mating system, growing environment and time (focused on time points 1 and 3) using a linear mixed model as described previously. After correcting for multiple testing, we identified metabolites showing both a significant 3-way interaction involving mating system, and a strong response to the common garden environment (through fold-change comparisons).

## RESULTS

### Natural population inbreeding does not reduce fitness in a novel common garden environment

Of the 310 transplanted individuals, 251 (79.0%) survived the first winter, with no significant effect of either mating system (Table 1: Likelihood Ratio statistic (LR-stat = 0.50, df = 1, P = 0.479) or initial rosette size (LR-stat = 0.91, df = 1, P = 0.340) on survival. Rosette size itself did not significantly vary with mating system (Fig 1b; LR-stat = 0.025, df = 1, P = 0.874), despite showing significant variation among populations (Fig 1b; LR-stat = 124.8, df = 12, P < 0.0001). Survival was markedly lower over the second winter, with only 34 individuals (11.0%) surviving to spring 2014, and again no effect of mating system (Table 1; LR-stat = 1.76, df = 1, P = 0.185), although outcrossing populations showed evidence for being more resilient to the challenges of overwintering. Specifically, over the second winter less than 5% of individuals survived in four of the five inbreeding populations, whereas only two of the eight outcrossing populations showed such low mortality (Table 1). The overall low survival over the second winter may in part be due to the milder winter temperatures experienced over the winter of 2013 (average 2.44°C) compared to 2012 (average 4.04°C; Fig S2). Despite no significant effect of inbreeding, the fixed effect of population explained 17.2% of variance in survival over the second winter (LR-stat = 36.8, df = 12, P = 0.0002), but was not significant for survival over the first winter. Interestingly, the inbreeding populations showed both the highest (PTP) and lowest (LPT, TC, KTT) rates of survival over the second winter, which emphasises population is more important than mating system for explaining patterns of survival.

**Table 1:** The proportion of surviving plants per population in the common garden on 28^th^ May 2013 and a year later on 21^st^ May 2014.

| Site<sup>a</sup> | Decimal degrees<sup>b</sup> |  | N<sup>c</sup> | Proportion surviving |  |
| --- | --- | --- | --- | --- | --- |
|  | Latitude | Longitude |  | 2013 | 2014 |
| SAK | 42.7044 | 86.2086 | 20 | 0.700 | 0.200 |
| LPT | 42.5797 | 80.3875 | 20 | 0.700 | 0.000 |
| SBD | 44.9389 | 85.8703 | 20 | 0.750 | 0.300 |
| MAN | 45.6703 | 82.2753 | 20 | 0.750 | 0.250 |
| TC | 45.2417 | 81.5175 | 20 | 0.750 | 0.000 |
| IND | 41.6214 | 87.2122 | 40 | 0.775 | 0.100 |
| TSS | 45.1925 | 81.5839 | 20 | 0.800 | 0.050 |
| TSSA | 45.1908 | 81.5906 | 20 | 0.800 | 0.050 |
| RON | 42.2614 | 81.8464 | 39 | 0.821 | 0.051 |
| PIN | 43.2689 | 81.8314 | 39 | 0.846 | 0.077 |
| KTT | 41.6206 | 83.7875 | 18 | 0.944 | 0.000 |
| PCR | 44.0042 | 83.0739 | 20 | 0.950 | 0.100 |
| PTP | 41.9278 | 82.5142 | 14 | 1.000 | 0.429 |
| Outcrossing |  |  | 199 | 0.799 | 0.131 |
| Selfing |  |  | 111 | 0.806 | 0.114 |

<sup>a</sup> sites are ordered by increasing proportion of individuals surviving in 2013; shaded populations represent outcrossing populations and unshaded inbreeding populations.
<sup>b</sup> coordinates in decimal degrees describing the latitude and longitude of each study location
<sup>c</sup> number of individuals at the start of the experiment in November 2012.

Population mating system also did not affect flowering phenology, with 88% of plants flowering within a 20-day time period in May. However, there were clear population effects, with individuals from SBD (outcrossing) flowering earliest and those from KTT (inbreeding) flowering latest (Fig 1c). On 10^th^ May 2013, when just over 50% of plants were flowering, there was no effect of population mating system on the propensity of individual plants to flower (LR-stat = 1.87, df = 1, P = 0.171), but there were significant population-level effects (LR-stat = 89.91, df = 12, P < 0.0001), and also a small, significant positive effect of rosette size when transplanted on the likelihood of flowering (LR-stat = 4.20, df = 1, P = 0.041).

In mid-July, no measure of reproductive investment varied with respect to mating system (number of fruits: LR-stat = 3.00, df = 1, P = 0.083; average fruit length: LR-stat = 0.04, df = 1, P = 0.842), although significant differences between populations were apparent (number of fruits: LR-stat = 31.46, df = 12, P = 0.002, Length fruits: LR-stat = 52.49, df = 12, P < 0.0001; Fig S3a,b). The percentage of within-individual variance in fruit length was 36.6%, which is similar to that explained by maternal plant (34.5%) and population (28.9%). By contrast, in this analysis mating system explained ~0% variance in fruit length. The combined measure of relative reproductive output revealed no significant effects of mating system on relative reproductive effort at this single time point when controlling for the random effect of population (Fig 1d; LR-stat = 3.08, df = 1, P = 0.079). There was also no significant difference among populations in relative reproductive effort, when population was considered as a fixed effect for the analyses (LR-stat = 7.50, df = 12, P = 0.377).

While on average selfing populations showed lower reproductive output than outcrossing populations in terms of numbers of fruits per individual (Table 2), this was driven by very low values for one of the selfing populations (LPT), in contrast to another selfing population on Lake Erie (RON), which showed a reproductive output on the higher end of values observed in the outcrossing populations.

**Table 2:** Fitness-related trait means, sample sizes and 95% confidence intervals for the study populations, as well as when separated into inbreeding and outcrossing groups. Rosette area (estimated area of a circle in cm^2^ is given) was measured for all plants, whereas fruit length (based on an average of 5 fruits per plant), the total number of fruits at a mid-season timepoint and total reproductive output (fruit length*number of fruits) was measured for a subset of plants (as described in the main text).

| Population | N | Rosette area (cm<sup>2</sup>) |  | N | Average fruit length (mm) |  | Number of fruits |  | Total reproductive effort |  |
| --- | --- | --- | --- | --- | --- | --- | --- | --- | --- | --- |
|  |  | Mean | 95% CI |  | Mean | 95% CI | Mean | 95% CI | Mean | 95% CI |
| IND | 40 | 35.23 | [31.55; 38.90] | 12 | 25.85 | [22.60; 29.11] | 63.75 | [31.68; 95.82] | 1665.74 | [866.23; 2465.25] |
| PCR | 20 | 36.39 | [31.69; 41.09] | 6 | 25.09 | [22.04; 28.13] | 60.83 | [29.78; 91.89] | 1538.96 | [685.55; 2392.36] |
| SBD | 20 | 34.49 | [27.99; 40.99] | 6 | 26.9 | [23.68; 30.13] | 66.17 | [6.78; 125.56] | 1847.51 | [83.11; 3611.90] |
| TSS | 20 | 30.5 | [26.39; 34.60] | 6 | 18.75 | [13.29; 24.20] | 62.42 | [24.60; 100.23] | 1186.1 | [347.14; 2025.05] |
| SAK | 20 | 35.97 | [31.07; 40.87] | 6 | 30.13 | [24.64; 35.62] | 32.92 | [12.95; 52.88] | 966.36 | [400.70; 1532.02] |
| PIN | 39 | 28.83 | [25.91; 31.76] | 12 | 23.69 | [21.08; 26.29] | 56.08 | [29.06; 83.11] | 1419.48 | [620.00; 2218.96] |
| MAN | 20 | 36.58 | [31.73; 41.42] | 6 | 22.59 | [19.07; 26.12] | 49.42 | [31.76; 67.07] | 1147.84 | [598.50; 1697.19] |
| TSSA | 20 | 33.43 | [27.92; 38.94] | 6 | 17.94 | [8.75; 27.12] | 25.83 | [9.68; 41.99] | 550.8 | [-31.42; 1133.02] |
| **Outcrossing** | **199** | **33.57** | **[32.04; 35.10]** | **54** | **24.73** | **[23.34; 26.11]** | **56.82** | **[46.01; 67.64]** | **1428.58** | **[1132.89; 1724.27]** |
| KTT | 18 | 14.51 | [11.34; 17.67] | 6 | 27.27 | [25.26; 29.27] | 36.67 | [12.62; 60.71] | 1030.89 | [300.22; 1761.57] |
| RON | 39 | 34.53 | [30.64; 38.43] | 12 | 22.65 | [20.27; 25.04] | 64.71 | [31.00; 98.41] | 1559.56 | [693.16; 2425.96] |
| TC | 20 | 48.42 | [40.78; 56.06] | 6 | 25.68 | [22.01; 29.35] | 35.5 | [15.34; 55.66] | 1003.76 | [609.44; 1398.09] |
| LPT | 20 | 41.94 | [37.49; 46.39] | 6 | 33.45 | [25.76; 41.14] | 14.08 | [9.68; 18.49] | 544.68 | [373.39; 715.96] |
| PTP | 14 | 26.84 | [22.08; 31.60] | 4 | 24.62 | [22.08; 27.16] | 51.5 | [14.83; 88.17] | 1257.18 | [440.70; 2073.66] |
| **Selfing** | **111** | **34.15** | **[31.22; 37.09]** | **40** | **24.61** | **[22.68; 26.53]** | **41.38** | **[29.75; 53.00]** | **1107.79** | **[792.28; 1423.29]** |

Rosette area before transplantation showed a significant interaction between mating system and latitude (Fig S4a), with a significantly increase in size with latitude for inbreeding, but not outcrossing populations. Flowering time also showed a significant interaction between mating system and latitude (Fig S4c), with an increase in flowering at the early time point for outcrossing populations with latitude but a very low flowering rate in the selfing population found at the highest latitude (TC). There was also a weakly negative effect of latitude on mean fruit length, but the pattern was the same in outcrossing and selfing populations (Fig S4e). After controlling for multiple testing, survival rates in both 2013 and 2014, the number of fruits produced and relative reproductive output at a mid-season timepoint showed neither a significant effect of latitude, nor an interaction of latitude with mating system (Fig S4b, d, f).

### Physiological responses to novel environments are driven by time and experimental treatments, with limited impact of inbreeding

Plants growing outside for the metabolomics study were exposed to herbivores and disease during the time course of sampling, and we avoided sampling any heavily damaged or infected plants for metabolomics analysis. At time point 2 (7^th^ August), the earliest time of sampling leaves in the field, there was no evidence for herbivore damage or disease on any plants used in the metabolomics analysis. At time point 3 (on 5^th^ Sept), eight of the 18 plants analysed showed no herbivore damage, and five of the 18 plants showed damage to just 1 leaf. The remaining five plants showed 2 or 3 damaged leaves). Molluscs were most likely causing this minor damage, and damage was mostly restricted to older leaves, which were not sampled in our study (JB personal observation). Additionally, three individuals showed early signs of *Albugo candida* infection (a common oomycete parasite of the Brassicaceae). These three individuals included one inbreeding (RON) and two outcrossing individuals (PIN and TSS). Therefore, the observed rates of infection and low levels of herbivore pressure should not impact the observed absence of inbreeding effects on the metabolome.

At time point 3 (one month after transplantation) there was clear divergence between plants in the growth chamber and common garden in growth and appearance (Fig S5), which made this time point the most informative for examining inbreeding effects on physiological plasticity. The first five principal components (PCs) extracted from all compounds explained 50.1% variation in the metabolome (Table 3).

**Table 3:** The significance of population mating system (inbreeding or outcrossing), time, experimental environment and their interactions for explaining variance in the first five principal components generated using data from 936 metabolites.

| Principal component | % variance in metabolites | Final model (explanatory factors)<sup>a</sup> | Significance of final model<sup>b</sup> |
| --- | --- | --- | --- |
| PC1 | 19.86 | environment * time | LR-stat = 77.75, df=2, p<0.0001 |
| PC2 | 13.07 | time | LR-stat=42.80, df=2, p<0.0001 |
| PC3 | 7.00 | environment * time * mating system | LR-stat=7.33, df=2, p=0.026 |
| PC4 | 5.29 | environment * time | LR-stat=19.99, df=2, p<0.0001 |
| PC5 | 3.08 | environment * time * mating system | LR-stat=7.06, df=2, p=0.029 |

<sup>a</sup> minimal adequate linear mixed model to explain variation in the principal component, generated by removing factors in order of complexity from a starting model of a 3-way interaction between all three fixed factors.
<sup>b</sup> likelihood ratio test statistic, associated degrees of freedom and p-value on comparison of the final model to the next most simple (nested) model.

Plotting PC1 (19.9% variance) against PC2 (13.1%) showed clear evidence for divergence in metabolite profiles at time point 3 compared to the earlier time points, particularly in the outdoor growth conditions, but with no strong effects of mating system (Fig 2a). Variation in these five PCs were mostly influenced by the interacting effects of growing environment and time, rather than mating system (Table 3). At time points 1, there was some clustering by population in both the growth cabinet and common garden environment, whereas at time point 3 this was only clear in the common garden, but in neither case was the pattern strongly associated with mating system or latitude (Fig S6). Time was a significant fixed factor for all five PCs and growing environment-by-time interactions were significant for four of the PCs. Despite a strong metabolomic shift over time in the experiment, there was no evidence for altered physiological plasticity at time point 3 in inbreeding populations. The direction and magnitude of change in PC1 and PC2 were mostly consistent across families and independent of population mating system (Fig 2b-2e). PC3 to PC5 (Fig S7a-f) also showed no effect of mating system on metabolomic plasticity, except that inbreeding populations showed greater variance in the magnitude of the metabolomic shift for PC5 (Fig S7f).

**Figure 2:**
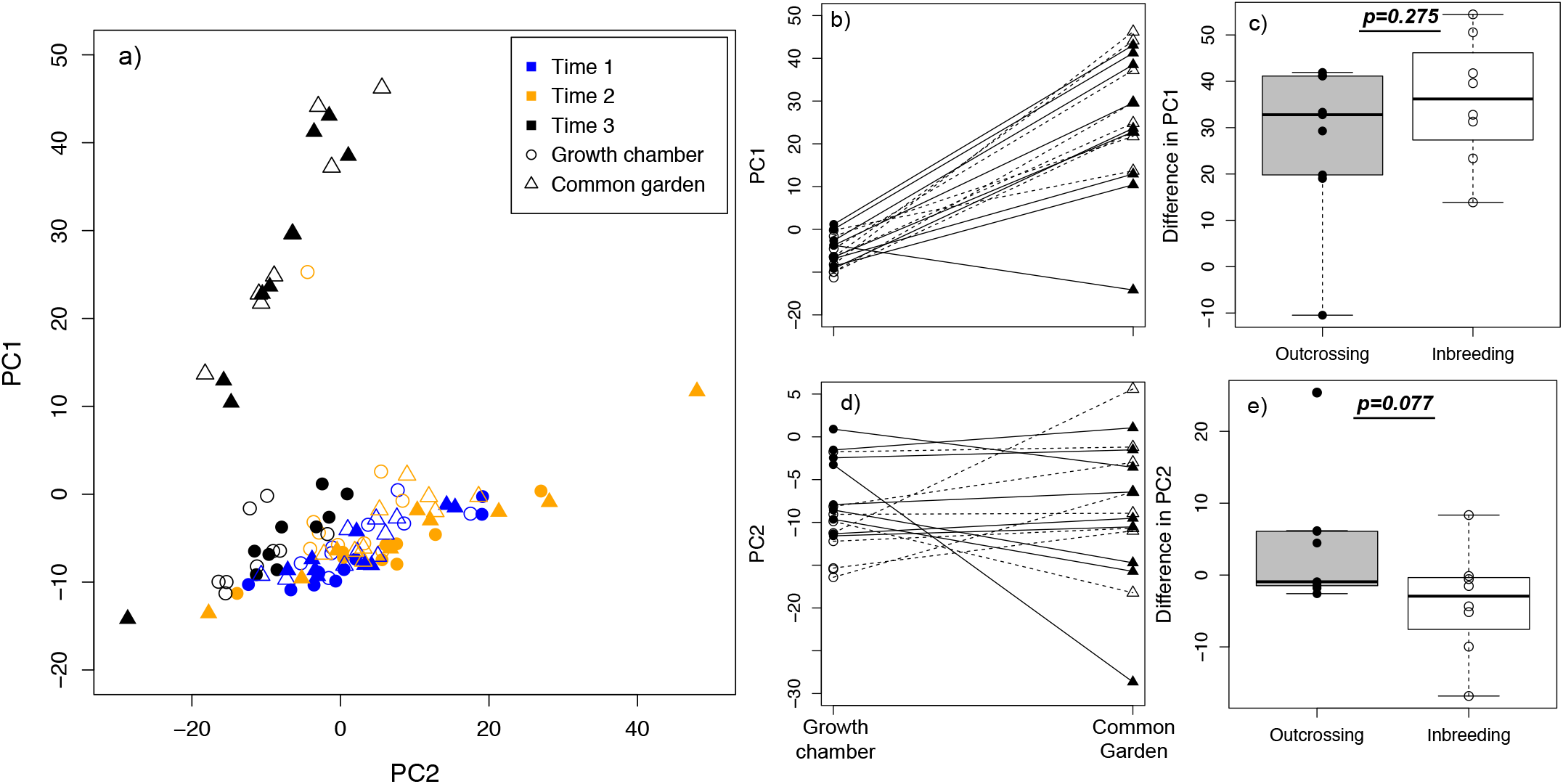
Multivariate variation and plasticity in the *Arabidopsis lyrata* metabolome with respect to time, mating system and experimental treatment: (a) visualised by plotting principal components 1 and 2, with colours indicating three different time points of sample collection (see key, Time 1= pre-transplant, Time 2=1 day after transplant, Time 3 = 1 month after transplant), symbols indicating environments (circles = growth chamber, triangles = common garden) and fill denoting mating system (open= inbreeding; filled = outcrossing); (b,d) plots of PC1 and PC2 values, respectively, for each individual at time point 3 for the two environments, with lines joining related individuals from the same family. Dashed lines (and open shapes) connect individuals from inbreeding populations and solid lines (and filled shapes) indicate those from outcrossing populations; (c,e) boxplots representing the change in PC1 and PC2 respectively between growing environments for related individuals grouped by population inbreeding status. The significance of population inbreeding on the magnitude of plasticity is indicated.

The diversity of metabolites changed significantly over time in the outdoor common garden environment, but again independent of mating system (Fig 3). Specifically, the metabolite richness (number of metabolites, when q = 0) showed a significant time*treatment interaction (P < 0.0001; Fig 3a) with fewer metabolites at time point 3 in the outdoor common garden, but not in the growth chamber. After accounting for relative abundance of metabolites (using q = 1 to reduce the contribution of rare compounds to estimated diversity), there was a significant time*treatment and significant mating system*treatment interaction (combined model significance: P < 0.0001). This was driven by a greater number of abundant compounds at time point 3 in the outdoor common garden, but also a tendency for inbred individuals to show an elevated number of abundant compounds relative to outcrossed populations at all time points in the growth chamber, but not in the common garden (Fig 3b).

**Figure 3:**
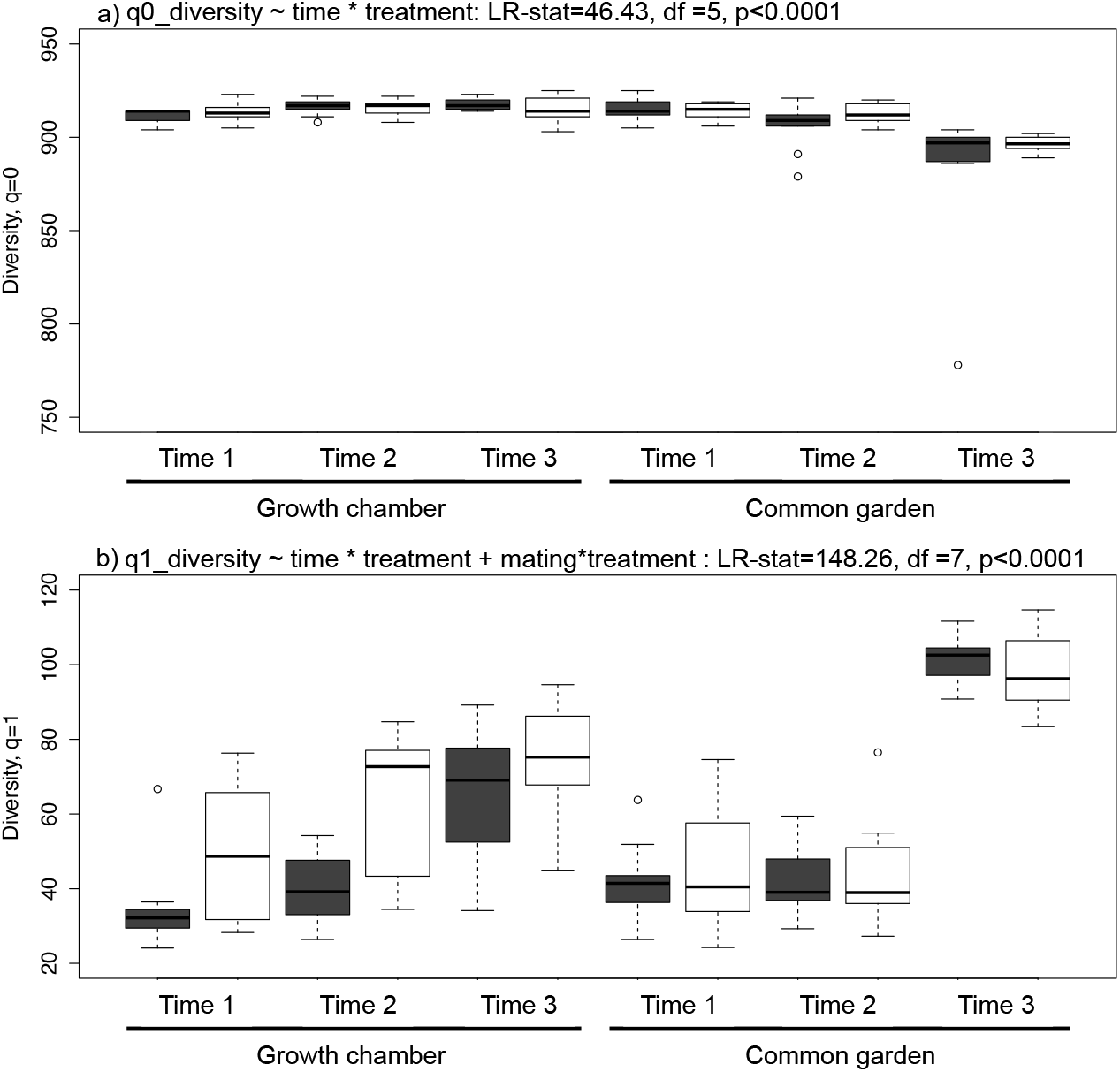
Metabolite diversity across the two growing environments for three different time points and individuals from outbred and inbred populations of *Arabidopsis lyrata.* Two measures of diversity are given based on (a) q=0 (analogous to number of metabolites observed) and (b) q=1 (reduced emphasis on metabolites at lower concentrations). Shaded boxes represent samples from outcrossing populations and open boxes those from inbreeding populations. The best model for explaining variation in diversity is given above each plot. The statistics represent the significant change in model likelihood when the full model was compared to the next simplest model.

### Individual metabolite data reveals limited effects of inbreeding, but clear changes in metabolites between growing environments over time

We found that 59.3% of the 936 metabolites showed significant (with a FDR of 5%) time*treatment interactions (Table 4a). Interactions involving mating system were only significant with 5% FDR for less than 0.5% metabolites (Table 4a). When the two growing environments (growth chamber and outdoor common garden) were analysed separately, time-by-mating system interactions were not significant for any metabolites following multiple testing correction (Table 4b). Similarly, no metabolites showed significant effects of just mating system in either growing environment (Table 4b). By contrast, the effect of time alone was significant for 36.0% (growth chamber) and 59.2% (common garden) of the metabolites (Table 4b). Notably, 1.6x more metabolites showed significant changes over time in the outdoor common garden compared to the growth chamber, which suggests that the outdoor environment was more physiologically challenging for the plant.

**Table 4:** Summary of results testing the effect of interactions between time, growing environment and mating system for 936 individual metabolites.

a)
| Interaction tested<sup>a</sup> | N p<0.05<sup>b</sup> | N p<0.01<sup>b</sup> | N (FDR 5%)<sup>b</sup> |
| --- | --- | --- | --- |
| 3-way | 125 | 48 | 7 |
| mating system*time | 76 | 22 | 4 |
| mating system *environment | 79 | 15 | 0 |
| time*environment | 601 | 496 | 555 |

<sup>a</sup> interactions between mating system (inbreeding or outcrossing), environment and time (time points 1 and 3 only)
<sup>b</sup> number of individual metabolites significant at different p-value thresholds, or with a false-discovery rate (Benjamini-Hochberg method) of 5%

b)
| Growing environment | Interaction tested<sup>a</sup> | N p<0.05<sup>b</sup> | N p<0.01<sup>b</sup> | N (FDR 5%)<sup>b</sup> |
| --- | --- | --- | --- | --- |
| Growth chamber | mating*time | 85 | 27 | 0 |
| Growth chamber | mating | 77 | 20 | 0 |
| Growth chamber | time | 405 | 305 | 337 |
| Common garden | mating*time | 42 | 5 | 0 |
| Common garden | mating | 63 | 15 | 0 |
| Common garden | time | 586 | 496 | 554 |

<sup>a</sup> mating system by time interactions separately for each growth treatment, compared between time points 1 and 3
<sup>b</sup> number of individual metabolites significant at different p-value thresholds, or with a false-discovery rate (Benjamini-Hochberg method) of 5%

Of the 106 confidently identified compounds, 27 were > 1-fold higher and 18 were > 1-fold lower in the outdoor common garden samples relative to the growth chamber samples at time point 3 (Table S3). Compounds that showed the strongest fold-changes included one annotated as the vitamin ascorbate (also annotated as D-Glucuronolactone), several members of the TCA cycle (S-malate, citrate and phosphoenolpyruvate) and several sugar phosphates associated with glycolysis and the pentose phosphate pathway (D-ribose 5-phosphate and D-glucose/D-Fructose 6-phosphate). The sugar sucrose was also 1.2-fold higher in the outdoor common garden samples. By contrast, 12 of 18 compounds that showed the greatest decrease in the common garden samples were amino acids or amino acid derivatives. Only three of the compounds showing > 1-fold change in the common garden samples also showed significant mating system effects at time point 3. For two of these compounds (dUMP and D-glucose/D-fructose-6-phospate), the effects of inbreeding were only apparent in the common garden sample set (Fig S8a,b). A significant negative effect of inbreeding in the common garden was only observed for the amino acid L-Asparagine (Fig S8c).

## DISCUSSION

In this study, we found that highly inbred and genetically depauperate populations of *A. lyrata* sampled from multiple genetic lineages (Foxe et al., 2010; Buckley et al., 2016) show similar fitness and short-term physiological responses to a change in environment as highly outcrossing and genetically diverse populations. Specifically, inbreeding populations showed similar survival rates and reproductive output to outbred populations over a two-year period in a common garden environment outside their native range, although relationships between several fitness traits and latitude of origin did vary between outcrossing and inbreeding populations. Instead, population-level effects, including latitude of origin, consistently explained more variation in fitness-related traits. Furthermore, we found that mating system had very little impact on the magnitude or plasticity of physiological responses to the novel environment, and only minor effects on metabolite diversity. Together, these results show that natural variation in inbreeding, and associated changes in genetic variation, do not negatively impact on short-term responses to changing environmental conditions.

### Natural population inbreeding does not reduce fitness in a novel common garden environment

Survival over the first winter was high in the common garden, but was much lower over the second winter, likely due to milder winter conditions resulting in root degradation over the winter of 2013/2014. Such mild winter conditions are rarely encountered in their native range around the North American Great Lakes region, where sub-freezing temperatures and snow cover are expected overwinter in all of the populations sampled. Variation in survival was not significantly explained by population mating system in either year, although populations did significantly differ in survival over the second winter, suggesting that factors other than mating system may be driving over-winter tolerance.

One hypothesis is that local adaptation to conditions in their native range might influence their rates of survival in this new environment. However, we found no evidence for an association between population latitude and survival in either year of this study. Three populations were sampled on the Bruce Peninsula on Georgian Bay (TSSA, TSS, TC) where the growing season is shorter than in the other populations sampled; previous surveys have found that plants typically flowered and seeds were produced approximately 1 month later in these northern populations than those growing on the shores of Lake Erie (RON, LPT, PTP) (personal observation). Although the Bruce Peninsula populations showed similar survival to one another, this was not true of the Lake Erie populations, suggesting that source latitude (and geographic proximity) is not a major driver of overwinter survival. In a different common garden experiment within the native range of *A. l. lyrata,* outcrossing populations showed higher survival rates after the first year, but over two years there was no effect of mating system on survival (Oakley, Spoelhof, and Schemske, 2015). Together with our results, these data suggest that reduced genetic variation due to inbreeding is not a consistent driver of variation in survival in this species.

A shift to selfing (and associated inbreeding) might also be associated with altered selection on growth rates, flowering traits and reproductive output (e.g. Sicard and Lenhard, 2011; Tedder et al., 2015). However, based on data from one mid-season time point, we observed no effect of mating system on the time to first flowering or measures of relative reproductive investment. Despite significant population effects on flowering time, the number of fruits produced and mean fruit length, the combined measure of relative reproductive output (a proxy for seed production) did not significantly vary among populations. By contrast, effects of mating system have been observed on flower and seed production traits in *A. l. lyrata* in two recent common garden experiments, conducted both in the native (Oakley *et al.* 2015) and non-native range (Willi *et al.* 2013), although effects were often only observed in one year and often varied between study years, supporting our finding of an absence of consistent effects of inbreeding on fitness. Furthermore, population-level, but not mating system effects, have also been observed for other flowering traits (including flower size and corolla length) in *A. l. lyrata* sampled from the same geographic region (Carleial, van Kleunen, and Stift, 2017b). These population effects may represent local adaptation to their native population environments, or simply phenotypic variation stochastically fixed in different regions following postglacial colonisation of the Great Lakes region.

Supporting the hypothesis of local adaptation shaping among-population trait variation, rosette area at time of transplant to the common garden tended to increase with latitude, but only in the selfing populations, which may reflect faster growth rates of individuals as an adaptation to shorter growing seasons. Furthermore, mean fruit length (correlated with the number of seeds in a fruit) declined with increasing latitude, which again may reflect an adaptation for more rapid reproduction under shorter growing seasons. However, the number of fruits did not vary with population latitude of origin, and this resulted in no significant variation in relative reproductive effort in this common environment for populations from different latitudes. Interestingly, flowering time was faster for outcrossing populations from higher latitudes, consistent with a faster transition to flowering with the more contracted growing season. Yet, the high latitude selfing population, TC, showed both a slower transition to flowering, and larger rosette size at transplantation, which could either be a consequence of its ability to self-fertilise, reducing its dependency on pollinators in a short growing season, or that it must reach a larger rosette size before it transitions to flowering. Nevertheless, we included just one selfing population at a higher latitude in our study, which was associated with a distinct rocky alvar habitat rather than sand dune habitat, so the effect of inbreeding on local adaptation is difficult to determine. Further sampling of high latitude inbreeding populations is therefore necessary to test whether populations with differing mating systems may diverge in their adaptive response to the broad environmental gradients represented by latitude.

The populations included in our study represented multiple genetic lineages, predicted to have colonised the Great Lakes region through different postglacial dispersal corridors (Hoebe, 2009; Foxe et al., 2010). However, our fitness data suggests that genetically (and often geographically) clustered populations are not more similar to each other in survival and flowering traits, than populations from distinct genetic clusters. For example, RON, LPT, PTP form a genetic cluster with PIN and PCR, but do not show more similarity in fitness-related traits than populations sampled from other genetic clusters (e.g. TC, TSS, TSSA).

The absence of strongly negative inbreeding effects on fitness in our common garden, even when plants were exposed to novel and stressful winter conditions is consistent with other studies, which have found that fitness differences between planting sites, experimental treatments (e.g. exposure to herbivory) or genetic families are frequently larger than the effects of experimental inbreeding (Ivey, Carr, and Eubanks, 2004; O’Halloran and Carr, 2010; Murren and Dudash, 2012). Our study also supports the hypothesis that North American populations of *A. l. lyrata* show only weak negative effects of natural inbreeding in most traits studied (Stift et al., 2013; Oakley, Spoelhof, and Schemske, 2015; Carleial, van Kleunen, and Stift, 2017a). The inbreeding populations used in this study have persisted following postglacial expansion into the Great lakes region (Hoebe, 2009; Foxe et al., 2010), so selection could have removed those individuals with the greatest inbreeding load. Alternatively, growing plants in a novel environment to which all populations are maladapted could have consistently reduced inbreeding depression effects in our study (as predicted by a theoretical study: Ronce et al., 2009). By contrast, experimentally-induced selfing has negatively affected germination rates and vegetative biomass (Carleial, van Kleunen, and Stift, 2017a), as well as reduced survival and reproductive effort (Willi, 2013; Oakley, Spoelhof, and Schemske, 2015) relative to experimentally-outbred progeny in both outcrossing and selfing populations of *A. l. lyrata.* Together, these data suggest that natural variation in inbreeding, in the absence of strong inbreeding depression, does not necessarily result in reduced fitness in a novel environment.

### Physiological responses are driven by time and growing environment, not history of inbreeding

Contrary to results from experimental-inbreeding studies (e.g. Kristensen et al., 2008; Campbell, Thaler, and Kessler, 2012), we found limited evidence that inbreeding altered physiological responses, as measured by shifts in the metabolome, to a novel common garden environment. We found no mating system effect on physiological plasticity through variation in the first two Principal Component (PC) axes, which together explained more than 30% of variation in the metabolome, and only minor effects of ! inbreeding on variation in PC3 and PC5, which together explained just 10.1% of variation ! in the metabolome. Instead, the first two PCs (explaining 30% of variation in the Į· metabolome) were strongly influenced by interactions between growing environment and ¡ time since transplantation. These results suggest that naturally inbred populations retain a similar physiological plasticity under different environmental conditions as outbred populations. Such a result contrasts with the effects of experimental inbreeding on î important biosynthetic pathways related to specific stressors, such as anti-herbivore defense induction (Campbell, Thaler, and Kessler, 2012; Kariyat et al., 2012). One explanation for the absence of mating system effects in our study is that the common garden treatment (one month growing in late summer) was not stressful enough to detect ! inbreeding effects on stress-related processes (Murren and Dudash, 2012; Schou, Kristensen, and Loeschcke, 2015). This suggests that testing physiological responses to known stressors in *A. lyrata*, for example herbivores or pathogens, might reveal physiological costs to inbreeding. However, experimental evidence using *A. lyrata* from) these same populations suggested no consistent negative effect of inbreeding on resistance to the pathogen *Albugo candida,* although the underlying physiological defense responses î were not measured (Hoebe et al., 2011). Furthermore, the induction of defenses by generalist herbivores in *A. lyrata* was unaffected by population inbreeding status (Joschinski, van Kleunen, and Stift, 2015). We found no significant effects of mating system on variation in amounts of individual metabolites in either environment, with ! significant mating system by time interactions only observed for a small number of ! compounds. Rather, changes over time in the common garden, relative to the growth Į· chamber, dominated the response of individual metabolites. Overall, the similarity of ¡ responses of individuals from inbreeding and outcrossing populations in our study suggests) that the reduced heterozygosity resulting from multiple generations of selfing has not compromised their ability to physiologically respond to contrasting environmental conditions.

The absence of effects of natural variation in mating system on plant physiology are perhaps expected, as the plant metabolome shows high plasticity under different growing environments (Brunetti et al., 2013). In our experiment, the plants grown in the outdoor common garden were exposed to a range of potential abiotic and biotic stressors, which are known to significantly alter the leaf metabolome (Sutter and Muller, 2011; Escobar-Bravo, Klinkhamer, and Leiss, 2017), often in ways specific to different stressors (Obata and Fernie, 2012). The observed changes in confidently-identified metabolites in our experiment are consistent with plants in the common garden responding to increased light intensity and levels of radiation, whereas plants in the growth chamber showed metabolic signatures of enhanced growth rates. Specifically, the common garden samples showed elevated levels of the vitamin ascorbate, a compound associated with UV-B tolerance (Wulff-Zottele et al., 2010; Kusano et al., 2011), as well as elevated levels of compounds linked to glycolysis (e.g. Glucose/Fructose −6-phosphate and Ribose-5-phosphate), and the TCA cycle (e.g. citrate, (S)-malate and succinate), suggesting elevated rates of photosynthesis (Wulff-Zottele et al., 2010). Conversely, the reduced levels of many important metabolites in growth chamber samples could reflect higher growth rates (e.g. Meyer et al., 2007), which is consistent with their higher rates of leaf production under controlled growth chamber conditions. Elevated growth rates in the growth chamber are also reflected in the elevated levels of amino acids in leaves in this growing environment, which is consistent with the higher levels of protein synthesis needed during growth (Hildebrandt et al., 2015). However, reduced amounts of amino acids in the common garden could also reflect resource constraints such as nitrogen availability (Obata and Fernie, 2012), although we used the same soil mix for both the common garden plot and growth chamber trays without additional fertiliser, so this seems unlikely to play a role in our experiment. Interestingly, statistical evidence for inbreeding effects on these highly responsive compounds in the common garden was limited to the amino acid, L-Asparagine. To understand the adaptive nature of these divergent responses to different growing environments, additional controlled experiments would be necessary to identify the key stressors driving observed physiological changes.

### Minor inbreeding effects on metabolite diversity under benign but not stressful conditions

Most metabolomic studies have used traditional multivariate approaches to interpret changes in metabolite composition (e.g. Davey et al., 2008; Kunin et al., 2009; Field and Lake, 2011). However, here we also estimated metabolite diversity using an approach based on changing the emphasis on the relative abundance across “species” using the parameter ‘q’ (Leinster and Cobbold, 2012). We found that inbreeding did not alter the total number of metabolites observed (equivalent to species richness, q = 0) across growing environments and time. However, when less emphasis was placed on low abundance metabolites (q = 1), inbred populations showed elevated metabolic diversity in the benign growth chamber environment relative to the common garden. This is consistent with inbreeding, specifically increased genome-wide homozygosity or exposure of deleterious alleles, having direct metabolic consequences (Pedersen, 1968; Cheptou and Donohue, 2011; Reed et al., 2012), and supports the importance of the environmental context of inbreeding depression (Kristensen et al., 2008; Bijlsma and Loeschcke, 2012).

Interestingly, experimentally inbred progeny from two self-incompatible *A. l. petraea* populations grown in a controlled environment also showed elevated expression of stress and photosynthesis related genes relative to outbred progeny (Menzel et al., 2015). Furthermore, inbred *Drosophila* lineages show changes to fundamental metabolic processes under both benign conditions and temperature stress (Pederson *et al.* 2008). It is therefore notable that mating system effects on metabolite diversity in our study were only observed in the constant growth chamber environment, and not in the outdoor common garden. However, the similarity of outcrossing and inbreeding populations in metabolite diversity in the common garden suggests that while inbreeding populations could suffer from the effects of reduced genome-wide heterozygosity, consistent with the observation of increased heterosis in these populations in other studies (Willi, 2013; Oakley, Spoelhof, and Schemske, 2015), this has limited effects on their short-term physiological capacity to respond to new environments. Instead, the metabolomic response to the environmental conditions experienced in our common garden overwhelms the relatively minor effects of genome-wide reduced heterozygosity in *A. lyrata.*

## CONCLUSIONS

Our results show that populations with a long history of inbreeding and associated reductions in genetic variation are not compromised in their ability to survive, reproduce and physiologically respond to a novel environment, at least relative to closely-related outcrossing populations. Such a finding could help to explain the general success of selfing lineages to colonise and adapt to a broad range of environments. The use of metabolomics to understand plastic physiological responses to novel environments and stressors offers promise for asking more specific questions linked to the understanding of the different type of pathways activated under a range of stressful conditions, as well as the impacts of genome-wide patterns of diversity on levels of metabolite diversity and plasticity. Together, these results offer new insights into the importance of intraspecific patterns of genetic variation for capacity to tolerate changing environmental conditions.

## ACKNOWLEDGEMENTS

We thank Glasgow Polyomics for generating and help analysing the metabolomics data, as well as the Scottish Centre for Ecology and the Natural Environment for providing space for the common garden and David Fettes for preparing the experimental site. We thank Natalie Hutchison who measured the length of fruits collected from the common garden. The comments of several anonymous reviewers have helped to improve the manuscript.

## AUTHOR CONTRIBUTIONS

JB and BKM designed the experiment. JB conducted the experiment, collected the data and performed the metabolite extractions. JB, CC, RD, KB and BKM analysed the metabolite data, and JB analysed the field data. JB and BKM wrote and revised the manuscript. Funding from a Natural Environment Research Council grant to BKM (NE/H021183/1)

## DATA ACCESSIBILITY STATEMENT

We will make all data available on an online repository, such as Data Dryad, should the manuscript be accepted for publication.

## SUPPORTING INFORMATION

**Table S1**: Location of populations used in this study, around the North American Great Lakes region, ordered by outcrossing rate (see Table S2).

**Table S2**: Details on the year of population sampling, population mating system variation, genetic diversity and heterozygosity and sample sizes for the experiments.

**Table S3**: List of 109 confidently identified compounds and their average log-fold changes between samples from the outdoor common garden and growth chamber at the initial pre-transplant time point and one month after transplant.

**Figure S1**: Layout of experimental common garden, illustrating how samples and populations were allocated to the plot.

**Figure S2**: Variation in temperature in the common garden field plot over the winter period (1^st^ Dec – 1^st^ March 2012 and 2013).

**Figure S3**: Summary plots of variation in reproductive investment among populations of *Arabidopsis lyrata* with respect to their mating system classification. Populations are ordered by outcrossing rates.

**Figure S4**: Regression plots illustrating the interaction between different fitness-related traits and latitude for the individuals from outcrossing and inbreeding populations.

**Figure S5**: Visual differences in *Arabidopsis lyrata* growing in the growth chamber and common garden one month after transplanting.

**Figure S6**: Principal component analysis of metabolite variation among sample populations at time points 1 and 3 for the growth chamber and outdoor common garden set of plants.

**Figure S7**: Plots representing plasticity in different principal components of metabolite variation in *Arabidopsis lyrata*.

**Figure S8**: Three compounds that show clear differences between the growth chamber and common garden at time point 3, as well as significant interactions between mating system and growing environment.

