## Supplementary tables and figures for "Changing environments and genetic variation: inbreeding does not compromise short-term physiological responses"

*American Journal of Botany* **Supporting Information**

The following Supporting Information is available for this article:

**Methods S1:** See main manuscript for all methodology

**SUPPLEMENTARY TABLES**

**Table S1**: Location of populations used in this study, around the North American Great Lakes region, ordered from highest to lowest outcrossing rate (see Table S2).

| Population | Lake | State/Province, Country | Population coordinates | | Location |
|---|---|---|---|---|---|
| | | | Latitude | Longitude | |
| IND | Michigan | Indiana, USA | N 41°37'17" | W 87°12'44" | Indiana Dunes National Lakeshore |
| PCR | Huron | Michigan, USA | N 44°00'15" | W 83°04'26" | Port Crescent State Park |
| SBD | Michigan | Michigan, USA | N 44°56'20" | W 85°52'13" | Sleeping Bear Dunes National Lakeshore |
| TSS | Huron | Ontario, Canada | N 45°11'33" | W 81°35'02" | Tobermory Singing Sands, Bruce Peninsula National Park |
| SAK | Michigan | Michigan, USA | N 42°42'16" | W 86°12'31" | Saugatak Dunes State Park |
| PIN | Huron | Ontario, Canada | N 43°16'08" | W 81°49'53" | Pinery Provincial Park |
| MAN | Georgian Bay | Ontario, Canada | N 45°40'13" | W 82°16'31" | Manitoulin Island |
| TSSA | Huron | Ontario, Canada | N 45°11'27" | W 81°35'26" | Tobermory Singing Sands Alvar, Bruce Peninsula National Park |
| KTT | - | Ohio, USA | N 41°37'14" | W 83°47'15" | Kitty Todd State Nature Preserve |
| RON | Erie | Ontario, Canada | N 42°15'41" | W 81°50'47" | Rondeau Provincial Park |
| TC | Georgian Bay | Ontario, Canada | N 45°14'30" | W 81°31'03" | Tobermory cliffs, Bruce Peninsula National Park |
| LPT | Erie | Ontario, Canada | N 42°34'47 " | W 80°23'15" | Long Point Provincial Park |
| PTP | Erie | Ontario, Canada | N 41°55'40" | W 82°30'51" | Point Pelee National Park |

**Table S2**: Details on the year of population sampling, population mating system variation, genetic diversity and heterozygosity and sample sizes for the experiments. Populations are ordered from highest to lowest outcrossing rate.

| | | | **Data from Foxe *et al.* (2010) – nine microsatellites** [a] | | | **Data from Buckley *et al.* (2016) – 6721 RAD loci** [b] | | **Common garden number individuals** [c] | | | |
|---|---|---|---|---|---|---|---|---|---|---|---|
| **Population** | **Year collected** | **Mating system** | **Outcrossing rate, $T_m$** | **SC** | $H_o$ | $H_o$ | $\pi$ | **N (1st germ)** | **N ( 2nd germ)** | **Total N** | **Subset N** |
| IND | 2011 | Outcrossing | 0.99 | 0.14 | 0.43 | 0.075 | 0.0011 | 30 | 10 | 40 | 12 |
| PCR | 2011 | Outcrossing | 0.98 | 0 | 0.23 | 0.075 | 0.0010 | 17 | 3 | 20 | 6 |
| SBD | 2011 | Outcrossing | 0.94 | 0 | 0.42 | 0.088 | 0.0012 | 11 | 9 | 20 | 6 |
| TSS | 2011 | Outcrossing | 0.91 | 0 | 0.361 | 0.073 | 0.0010 | 17 | 3 | 20 | 6 |
| SAK | 2011 | Outcrossing | 0.9 | - | - | 0.078 | 0.0012 | 14 | 6 | 20 | 6 |
| PIN | 2011 | Outcrossing | 0.84 | 0 | 0.18 | 0.068 | 0.0009 | 36 | 3 | 39 | 12 |
| MAN | 2011 | Outcrossing | 0.83 | 0.13 | 0.22 | 0.061 | 0.0009 | 18 | 2 | 20 | 6 |
| TSSA | 2011 | Outcrossing | 0.41 | 0.5 | 0.097 | 0.067 | 0.0010 | 13 | 7 | 20 | 6 |
| KTT | 2007 | Selfing | 0.31 | 1 | 0 | 0.006 | 0.0002 | 13 | 5 | 18 | 6 |
| RON | 2011 | Selfing | 0.28 | 1 | 0.028 | 0.029 | 0.0005 | 30 | 9 | 39 | 12 |
| TC | 2012 | Selfing | 0.18 | 0.88 | 0.181 | 0.002 | 0.0000 | 18 | 2 | 20 | 6 |
| LPT | 2012 | Selfing | 0.13 | 1 | 0.069 | 0.006 | 0.0002 | 14 | 6 | 20 | 6 |
| PTP | 2011/2012 | Selfing | 0.09 | 1 | 0.069 | 0.004 | 0.0004 | 10 | 4 | 14 | 4 |

[a] average outcrossing rate estimated through progeny arrays (*Tm)*, and proportion of self-compatible plants (SC), as well as observed heterozygosity ($H_o$) as estimated by Foxe *et al.* (2010) using nine microsatellites.
[b] observed heterozygosity ($H_o$) and nucleotide diversity ($\pi$) as estimated using 6721 RAD-seq loci in Buckley *et al.* (2016)
[c] sample sizes for two rounds of germinations (the second to supplement families with low germination rates in the first round), the total number of plants used for estimating survival (Total N) and the subset used for measuring fruit production for reproductive fitness estimates (Subset N - fruits)

**Table S3**: List of 109 confidently identified compounds and their average log-fold changes between samples from the outdoor common garden and growth chamber at the initial pre-transplant time point and one month after transplant. Compounds showing an increase or decrease in the outdoor common garden (OG) samples relative to the growth chamber (GC) samples are highlighted in green and blue, respectively, with the darkness of shading indicating the relative strength of the log-fold change (logFC). Note that 13 identified compounds had identical retention times and could not be distinguished during peak integration, so are combined on this table (see ** next to names).

| | | **Early timepoint** | **Late timepoint** |
|---|---|---|---|
| **Name** | **Formula** | **logFC OG / GC** | **logFC OG / GC** |
| ascorbate / D-Glucuronolactone ** | C6H8O6 | 1.38 | 6.41 |
| D-ribose 5-phosphate | C5H11O8P | -0.22 | 3.89 |
| citrate | C6H8O7 | 0.43 | 2.98 |
| Phosphoenolpyruvate | C3H5O6P | 0 | 2.97 |
| (S)-Malate | C4H6O5 | -0.19 | 2.58 |
| 2/3-Phospho-D-glycerate** | C3H7O7P | 0 | 2.57 |
| D-Threose | C4H8O4 | -0.29 | 2.49 |
| succinate semialdehyde | C4H6O3 | -0.1 | 2.46 |
| L-Carnitine | C7H15NO3 | 0.01 | 2.38 |
| D-glucose 6-phosphate / D-Fructose 6-phosphate ** | C6H13O9P | 0.39 | 2.23 |
| Glycine | C2H5NO2 | -0.53 | 2.17 |
| Orthophosphate | H3O4P | 0.01 | 2.17 |
| AMP / dGMP ** | C10H14N5O7P | 0.12 | 1.91 |
| Pantothenate | C9H17NO5 | 0.05 | 1.58 |
| N-Acetyl-D-glucosamine | C8H15NO6 | -0.2 | 1.47 |
| Guanine | C5H5N5O | -0.13 | 1.44 |
| L-Rhamnose | C6H12O5 | 0.05 | 1.28 |
| (R)-2-Hydroxyglutarate | C5H8O5 | -0.39 | 1.27 |
| D-Gluconic acid | C6H12O7 | 0 | 1.23 |
| L-Tyrosine | C9H11NO3 | 0.17 | 1.22 |
| N-Acetylneuraminate | C11H19NO9 | 0.06 | 1.2 |
| sucrose | C12H22O11 | -0.05 | 1.2 |
| 2-Deoxy-D-glucose | C6H12O5 | -0.1 | 1.13 |
| Pyruvate | C3H4O3 | 0.28 | 1.12 |
| Gallate | C7H6O5 | -0.25 | 1.12 |
| L-Tryptophan | C11H12N2O2 | -0.18 | 1.1 |
| Maleic acid | C4H4O4 | 0.02 | 1.07 |
| D-Glucono-1,4-lactone | C6H10O6 | 0.03 | 1.02 |
| Adenosine | C10H13N5O4 | -0.22 | 0.96 |
| Imidazole-4-acetate | C5H6N2O2 | -0.02 | 0.92 |
| 2-Oxoglutarate | C5H6O5 | -0.2 | 0.84 |
| 6-Phospho-D-gluconate | C6H13O10P | 0.16 | 0.83 |
| Acetoacetate | C4H6O3 | 0.23 | 0.82 |
| L-Gulono-1,4-lactone | C6H10O6 | 0.15 | 0.78 |
| L-Phenylalanine | C9H11NO2 | 0.22 | 0.76 |
| (R)-Lactate | C3H6O3 | -0.59 | 0.74 |
| Orotate | C5H4N2O4 | -0.28 | 0.74 |

| | | Early timepoint | Late timepoint |
|---|---|---|---|
| **Name** | **Formula** | **logFC OG / GC** | **logFC OG / GC** |
| cytosine | C4H5N3O | 0.03 | 0.72 |
| D-Galactarate | C6H10O8 | 0.05 | 0.66 |
| cis-Aconitate / L-Dehydroascorbate ** | C6H6O6 | 0.26 | 0.64 |
| D-Glucosamine-6-phosphate | C6H14NO8P | 0.09 | 0.6 |
| HEPES | C8H18N2O4S | 0.05 | 0.59 |
| Guanosine | C10H13N5O5 | 0.18 | 0.56 |
| Thiamin | C12H16N4OS | -0.15 | 0.52 |
| riboflavin | C17H20N4O6 | -0.03 | 0.48 |
| 1,3-Diaminopropane | C3H10N2 | -0.38 | 0.48 |
| L-Histidine | C6H9N3O2 | 0.25 | 0.47 |
| (R)-3-Hydroxybutanoate | C4H8O3 | -0.03 | 0.47 |
| 5-Aminolevulinate | C5H9NO3 | -0.24 | 0.46 |
| 4-Trimethylammoniobutanoate | C7H15NO2 | -0.1 | 0.37 |
| D-glucose | C6H12O6 | 0.08 | 0.36 |
| N2-Acetyl-L-lysine | C8H16N2O3 | -0.02 | 0.3 |
| Ala-Gly | C5H10N2O3 | -0.17 | 0.3 |
| L-Leucine / L-isoleucine ** | C6H13NO2 | 0.15 | 0.29 |
| 4-Aminobutanoate | C4H9NO2 | 0.14 | 0.29 |
| L-Kynurenine | C10H12N2O3 | 0.09 | 0.28 |
| 4-Aminobenzoate | C7H7NO2 | -0.02 | 0.24 |
| Spermidine | C7H19N3 | 0.48 | 0.24 |
| L-Valine / Betaine ** | C5H11NO2 | 0.76 | 0.21 |
| thymine | C5H6N2O2 | 0.09 | 0.15 |
| Pyridoxal | C8H9NO3 | -0.03 | 0.12 |
| 1-Aminopropan-2-ol | C3H9NO | -0.09 | 0.12 |
| Acetylcholine | C7H15NO2 | -0.07 | 0.12 |
| Inosine | C10H12N4O5 | -0.16 | 0.09 |
| Spermine | C10H26N4 | 0.03 | 0.02 |
| D-Galacturonate | C6H10O7 | 0.1 | 0.02 |
| Orotidine | C10H12N2O8 | 0.15 | 0.01 |
| 5'-Methylthioadenosine | C11H15N5O3S | -0.15 | -0.02 |
| L-2,4-Diaminobutanoate | C4H10N2O2 | 0.04 | -0.02 |
| N-Acetylornithine | C7H14N2O3 | -0.13 | -0.03 |
| D-Fructose | C6H12O6 | 0.52 | -0.06 |
| Phthalate | C8H6O4 | 0.28 | -0.07 |
| Fumarate | C4H4O4 | -0.04 | -0.12 |
| Oxalate | C2H2O4 | 0.75 | -0.12 |
| 1-Aminocyclopropane-1-carboxylate | C4H7NO2 | -0.12 | -0.14 |
| Methylmalonate / Succinate ** | C4H6O4 | -0.02 | -0.14 |
| N-acetyl-L-glutamate | C7H11NO5 | -0.27 | -0.26 |
| agmatine | C5H14N4 | 0.55 | -0.32 |
| Choline phosphate | C5H14NO4P | 0.2 | -0.38 |
| Malonate | C3H4O4 | 0.32 | -0.4 |
| L-Glutamate / O-Acetyl-L-serine ** | C5H9NO4 | -0.1 | -0.44 |

| | | Early timepoint | Late timepoint |
|---|---|---|---|
| **Name** | **Formula** | **logFC OG / GC** | **logFC OG / GC** |
| L-2-Aminoadipate | C6H11NO4 | 0.01 | -0.45 |
| L-Proline | C5H9NO2 | 0.08 | -0.47 |
| L-Ornithine | C5H12N2O2 | -0.16 | -0.55 |
| Choline | C5H13NO | -0.12 | -0.56 |
| Adenine | C5H5N5 | 0.2 | -0.61 |
| L-Lysine | C6H14N2O2 | 0.26 | -0.61 |
| L-Noradrenaline / Pyridoxine ** | C8H11NO3 | -0.05 | -0.69 |
| trans-4-Hydroxy-L-proline | C5H9NO3 | 0.11 | -0.79 |
| sn-glycero-3-Phosphocholine | C8H20NO6P | -0.51 | -0.83 |
| S-Adenosyl-L-homocysteine | C14H20N6O5S | -0.2 | -0.92 |
| L-Serine | C3H7NO3 | -0.23 | -1.23 |
| L-Methionine | C5H11NO2S | -0.19 | -1.31 |
| beta-Alanine / L-Alanine ** | C3H7NO2 | -0.09 | -1.64 |
| 5-Oxoproline | C5H7NO3 | -0.04 | -1.73 |
| L-homoserine / L-Threonine ** | C4H9NO3 | -0.21 | -2.01 |
| N-Acetylglutamine | C7H12N2O4 | 0.37 | -2.01 |
| Cytidine | C9H13N3O5 | 0.09 | -2.08 |
| Methylcysteine | C4H9NO2S | -0.56 | -2.1 |
| Picolinic acid | C6H5NO2 | -0.06 | -2.35 |
| Isonicotinic acid / Nicotinate ** | C6H5NO2 | -0.28 | -2.37 |
| L-Citrulline | C6H13N3O3 | -0.37 | -2.4 |
| L-Glutamine | C5H10N2O3 | -0.13 | -2.44 |
| L-Aspartate | C4H7NO4 | -0.27 | -2.45 |
| N(pi)-Methyl-L-histidine | C7H11N3O2 | 0.66 | -2.8 |
| L-Arginine | C6H14N4O2 | 0.14 | -2.95 |
| dUMP | C9H13N2O8P | 0.38 | -3.46 |
| L-Cysteate | C3H7NO5S | 0 | -3.88 |
| L-Asparagine | C4H8N2O3 | -0.01 | -8.23 |

### SUPPLEMENTARY FIGURES

**Figure S1**: Layout of experimental common garden, illustrating how samples and populations were allocated to the plot. The top panel shows the four experimental blocks, and the second shows one block in detail, with 5 individuals per population (10 for IND, PIN and RON) distributed systematically across the trays. Dark shading indicates outcrossing populations and no shading indicates inbreeding. Finally, the bottom plot shows an example of how 20 individual plants are distributed across each tray.

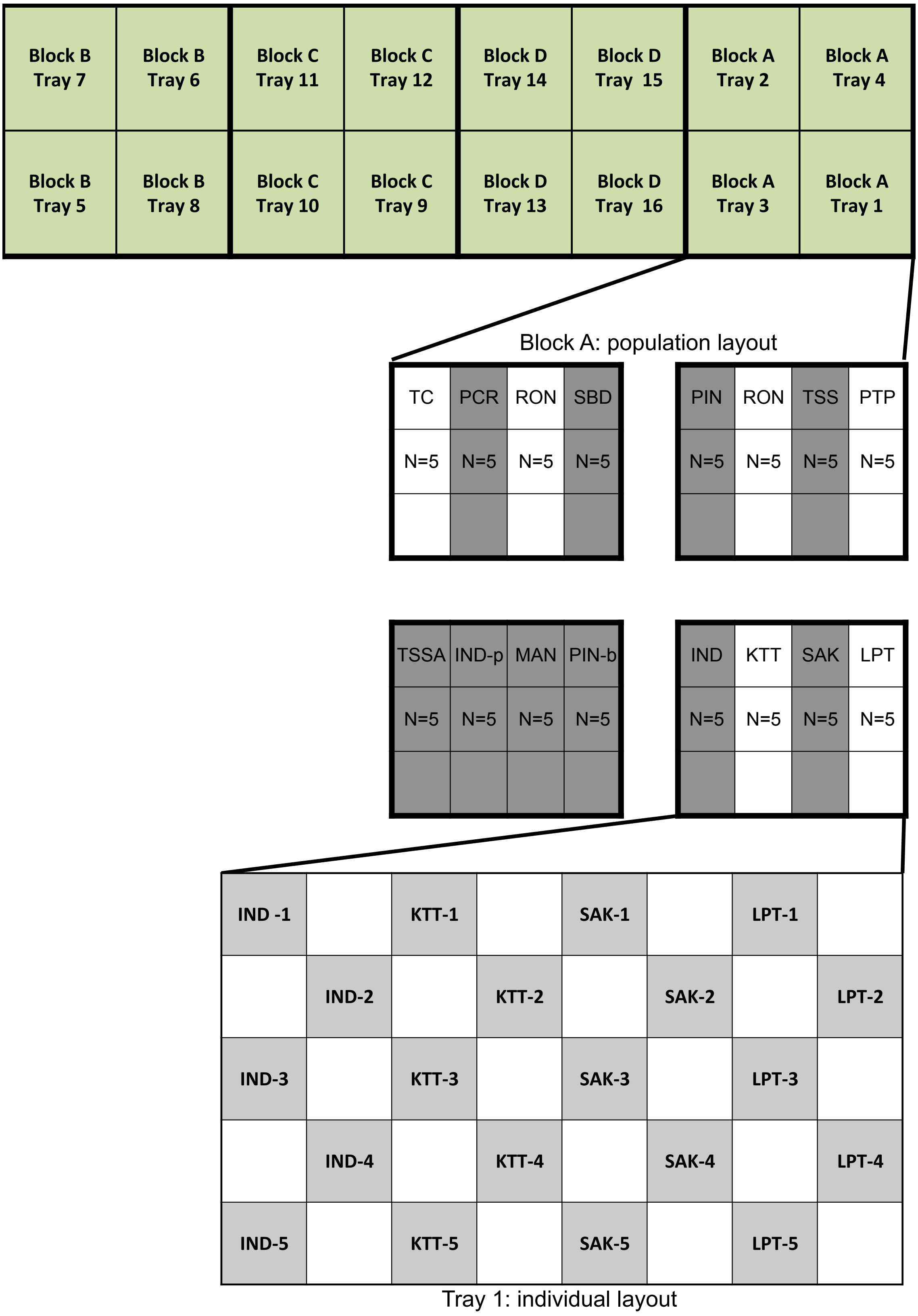

**Figure S2**: Variation in temperature in the common garden field plot over the winter period (1st Dec – 1st March 2012 and 2013). Temperature is averaged across two iButton dataloggers, one placed just under soil surface (soil temperature) and one just above the surface (air temperature). Points highlighted in green are those below 0°C, and show more days below this threshold in the winter of 2012 than the winter of 2013. These milder winter temperatures in Winter 2013 could have resulted in root degradation in plants, which normally grow under cold (freezing) winter conditions around the Great Lakes.

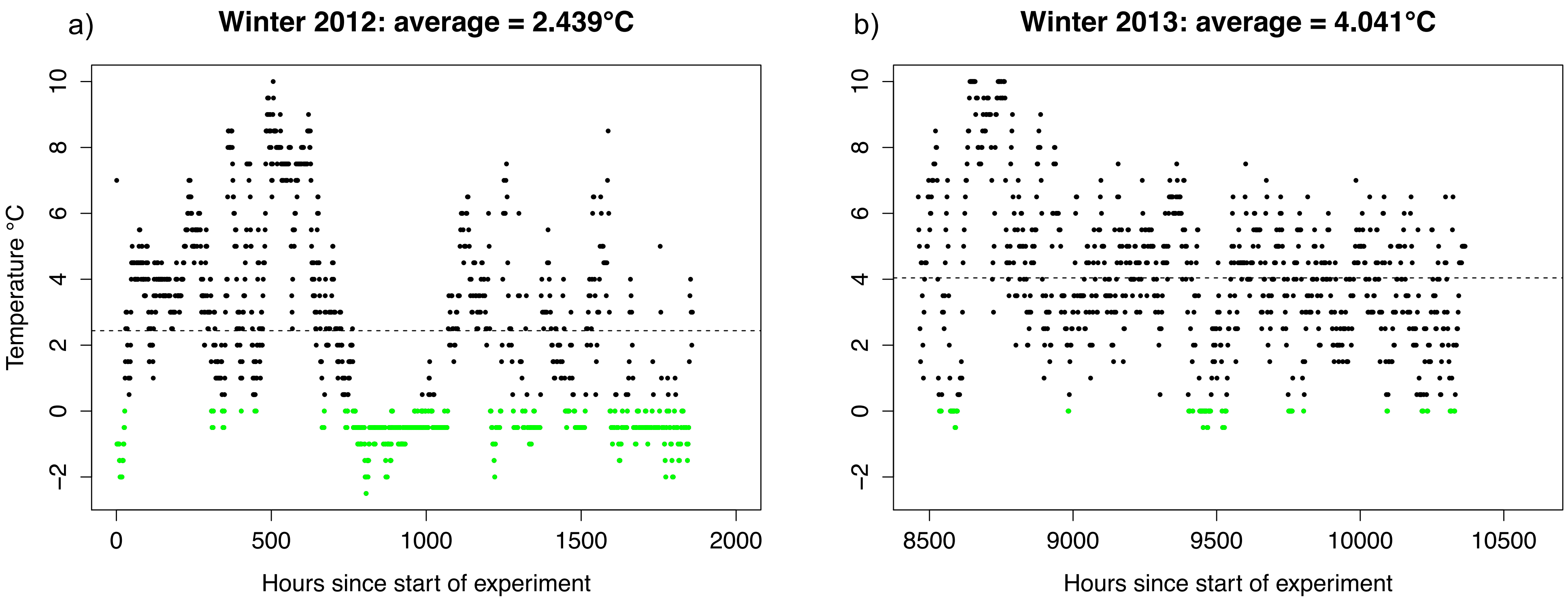

**Figure S3**: Summary plots of variation in reproductive investment among populations of *Arabidopsis lyrata* with respect to their mating system classification. (a) the number of fruits produced by a subset of individuals per populations as recorded on 1st July 2013; (b) the average length of fruits produced by that subset of plants per site. Dark grey boxes indicate outcrossing, and white boxes indicate inbreeding populations. Site codes and sample sizes are given below each box. Populations are ordered by outcrossing rates.

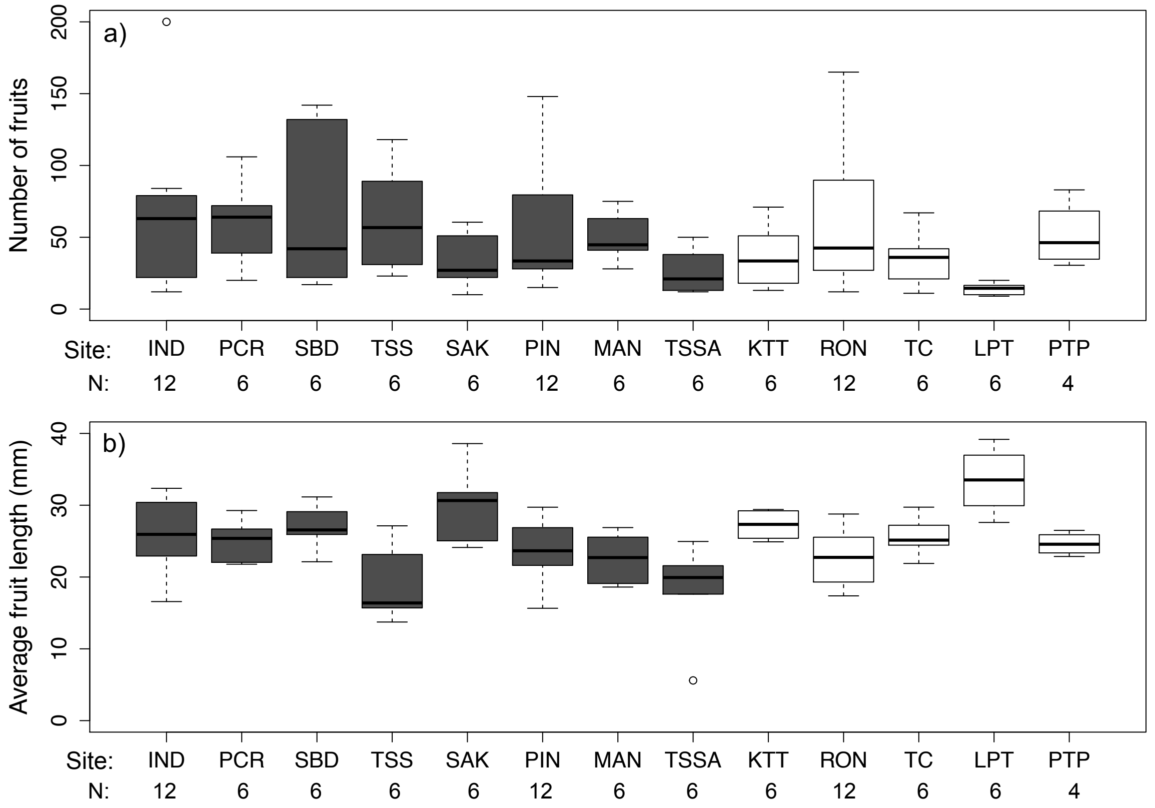

**Figure S4:** Regression plots illustrating the interaction between different fitness-related traits and latitude for the individuals from outcrossing (dark grey) and inbreeding (light grey) populations. (a) Rosette area in $mm^2$, (b) proportion of plants alive per population in spring 2014, (c) proportion of plants flowering per population at an early season timepoint (10th May 2013), (d) Number of fruits at a mid-season timepoint (1st July 2013), (e) mean fruit length per individual plant (mm), and (f) relative reproductive output at the mid-season timepoint (mean fruit length multiplied by number of fruits). The significance of the interaction between mating system and latitude, or latitude alone, is given where significant. Mating system alone was significant for none of the traits. With Bonferroni correction for multiple testing (p= 0.008 threshold), the interaction in (b) is not significant.

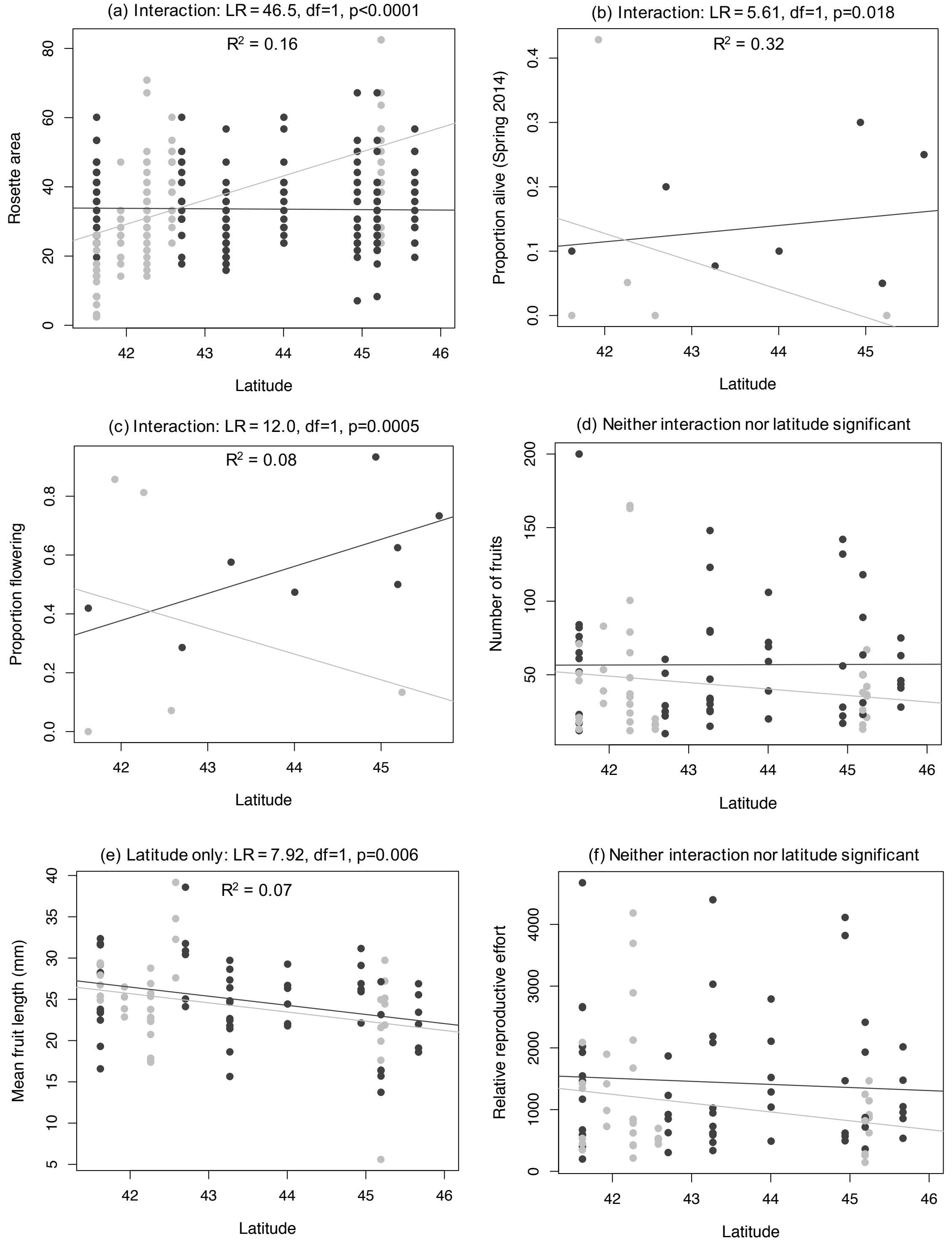

**Figure S5**: Visual differences in *Arabidopsis lyrata* growing in the growth chamber and common garden one month after transplanting. Photos representing the status of plants growing in (a,b) the growth chamber or (c,d) outside in the common garden at the time of transplanting to the common garden (time point 2, plots a, c) and after one month growing in those respective environments (time point 3, plots b, d). These photos illustrate the marked differences in growing environment experienced by the plants inside and outside, supporting the clear differences in metabolomic profile observed at time point 3, but not time point 2.

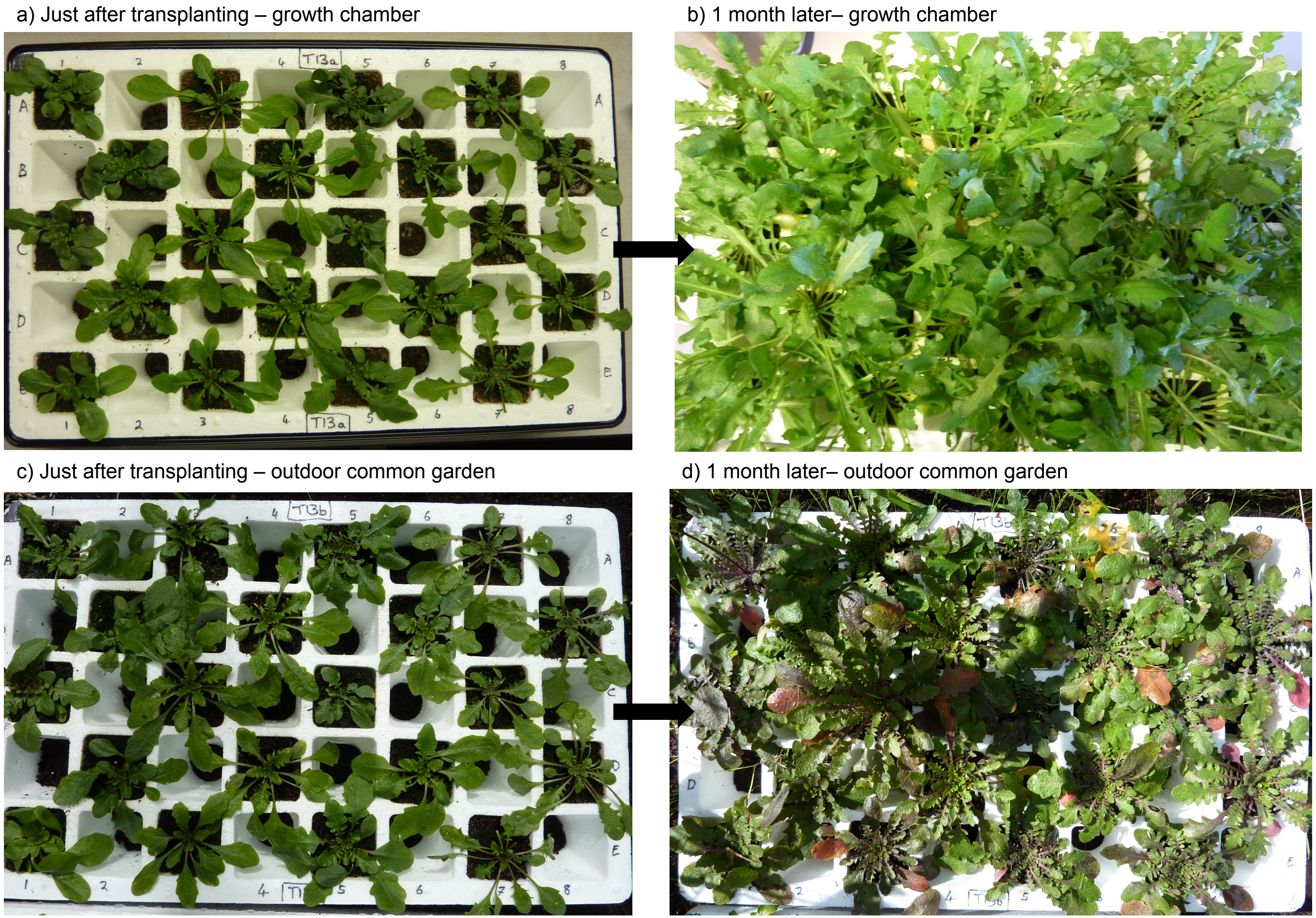

**Figure S6:** Principal component analysis of metabolite variation among sample populations at time points 1 and 3 for the growth chamber and outdoor common garden set of plants. Principal components 1 and 2 are plotted against each other for each set of samples with points coloured by population of origin (see colour key). (a) Time point 1 for the growth chamber set of plants, (b) Time point 1 for the outdoor common garden set of plants, (c) Time point 3 for growth chamber set of plants, and (d) Time point 3 for the outdoor common garden set of plants. These plots illustrate the minor effect of population in driving patterns of metabolomic variation.

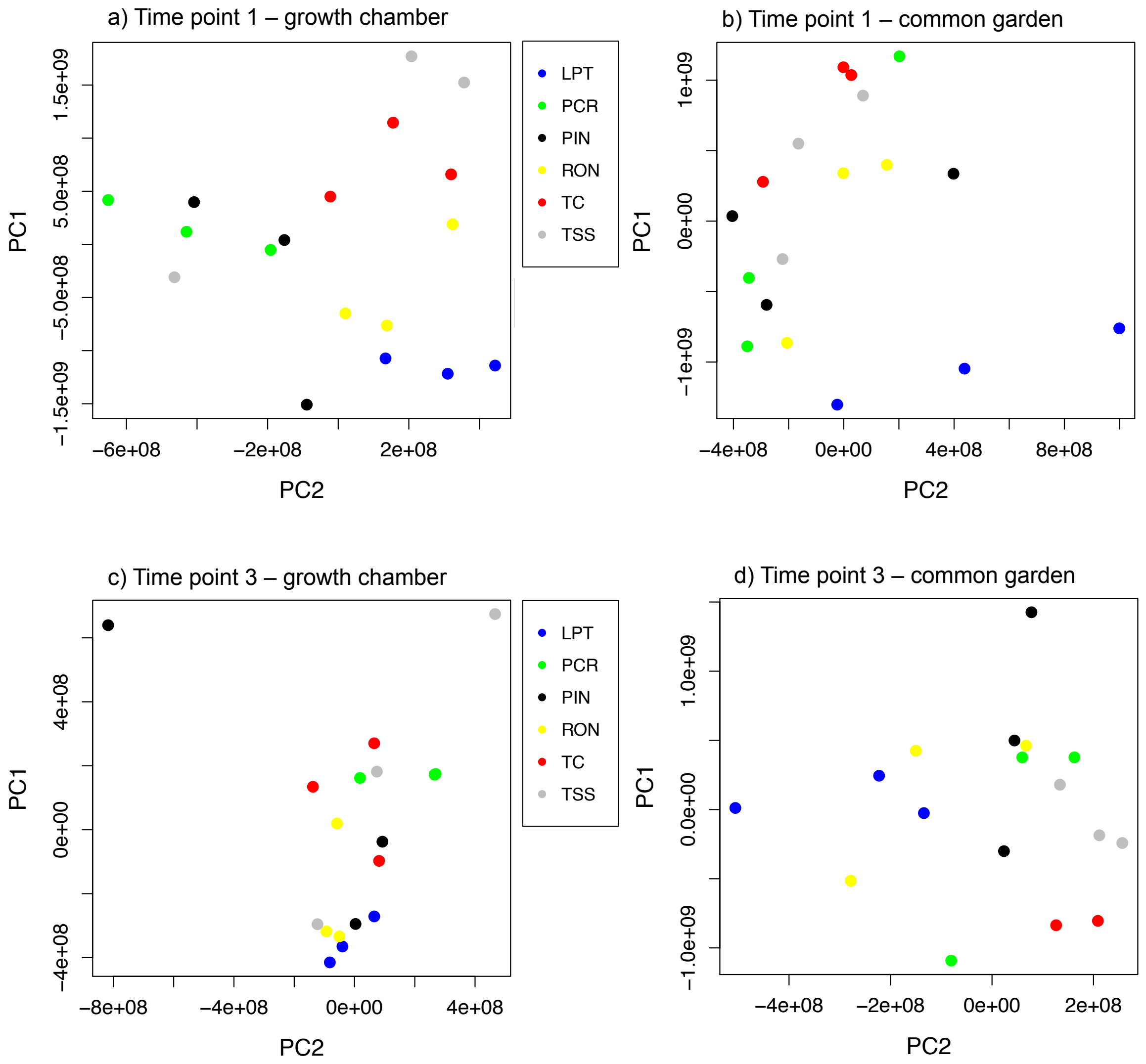

**Figure S7**: Plots representing plasticity in different principal components of metabolite variation in *Arabidopsis lyrata*. (a,c,e) represent plots of PC3, PC4 and PC5 values respectively for related individuals at time point 3 in the growth chamber and common garden. Lines connect individuals from the same family. Dashed lines (and open shapes) connect individuals from inbreeding populations and full lines (and filled shapes) connect those from outcrossing populations; (b,d,f) boxplots representing the change in each PC between related individuals in the growth chamber or common garden grouped by population inbreeding status. The significance of the difference in the magnitude of plasticity is given, as estimated by ANOVA with mating system as a fixed effect.

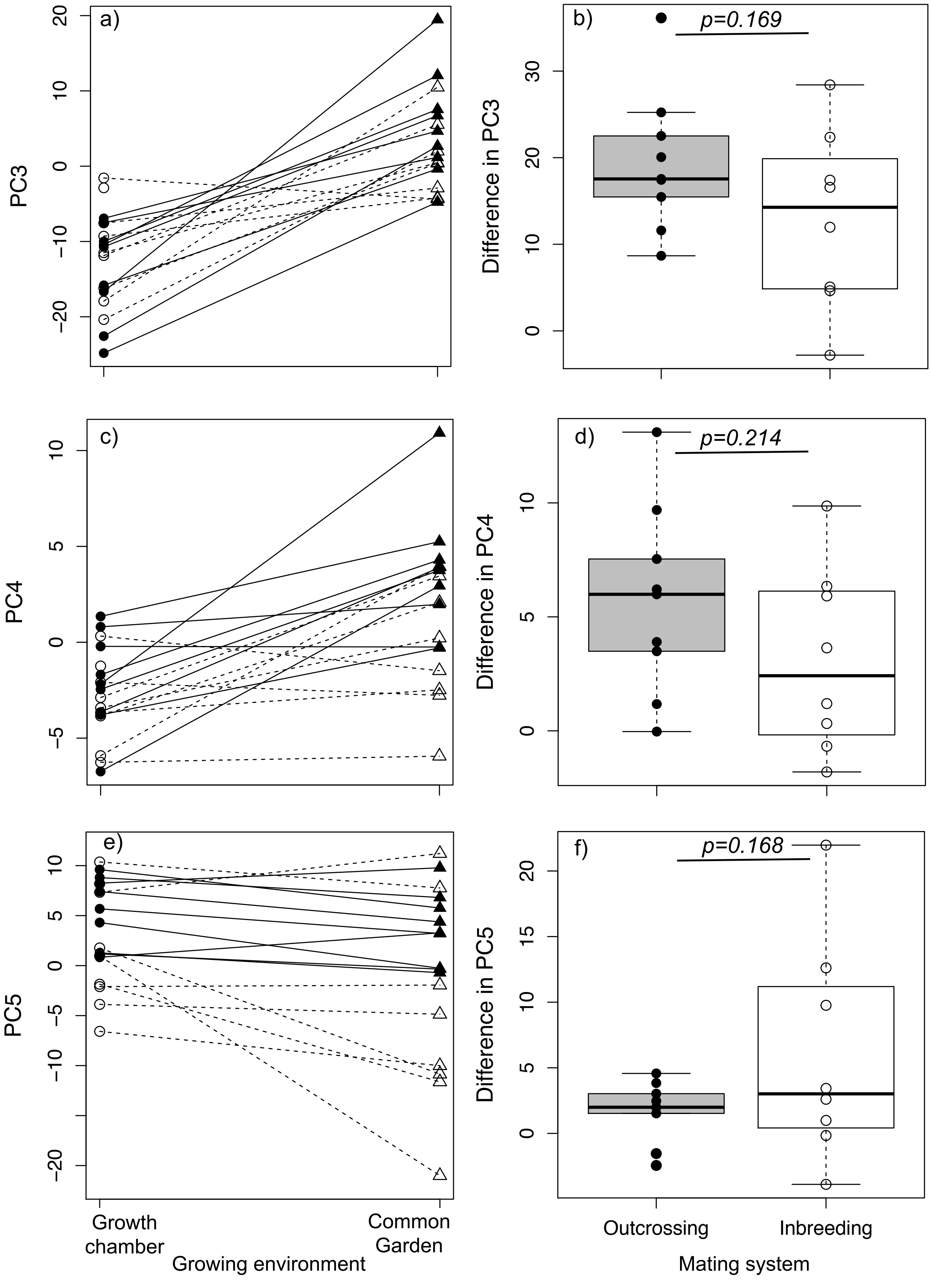

**Figure S8**: Three compounds that show clear differences between the growth chamber and common garden at time point 3 (see Table S3), as well as significant interactions between mating system and growing environment. The compounds are (a) dUMP, (b) D-glucose/fructose-6-phosphate (difficult to distinguish these metabolites with the given standards), and (c) the amino acid L-Asparagine. The y-axis represents the log2 values of the maximum peak height (peak intensity). Q-values represent the multiple-testing corrected p-values for the 3-way interaction between mating system, growing environment. Dark gray boxes represent outcrossing and open boxes represent inbreeding samples.

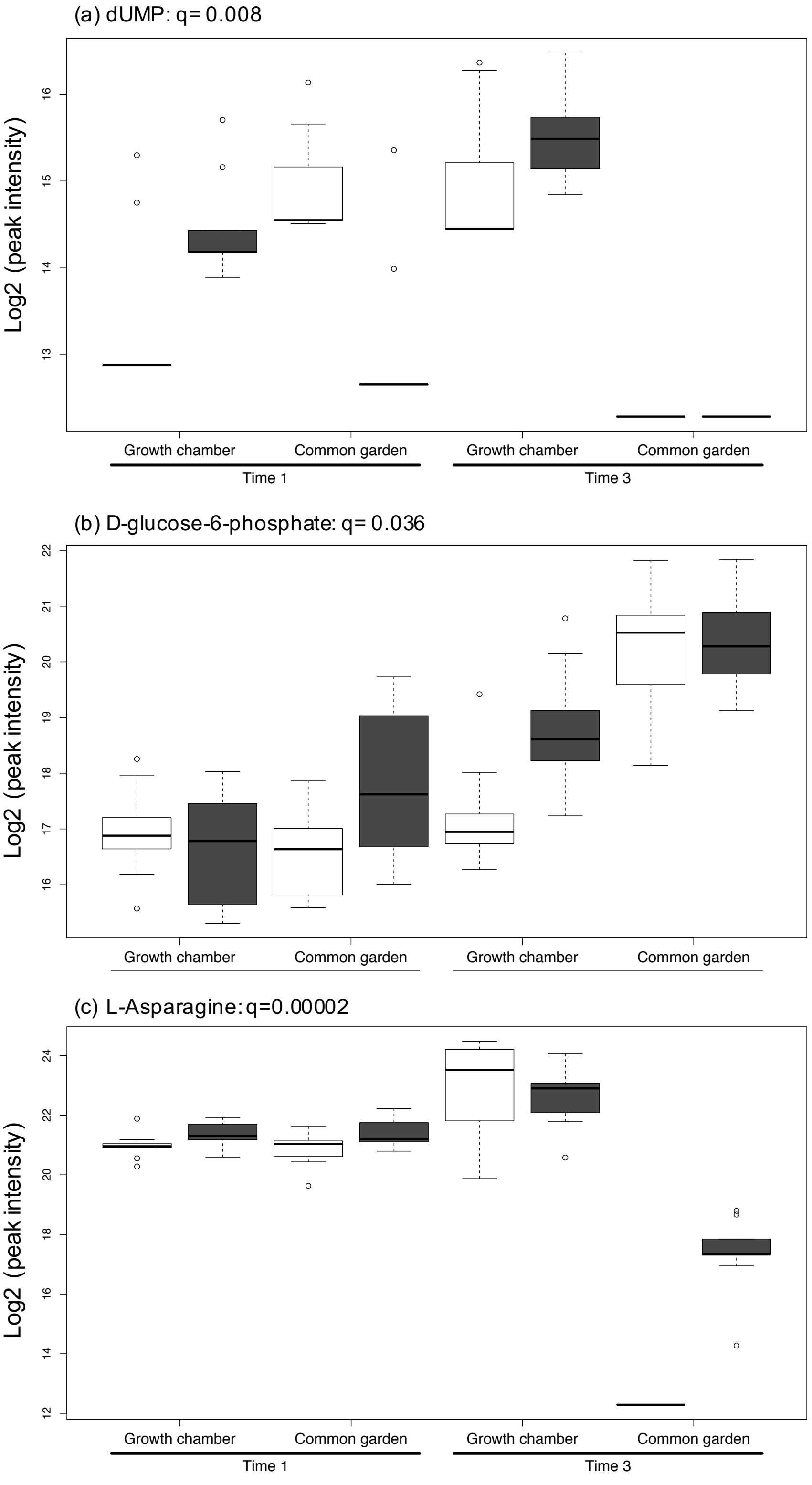
